# Characterization of Morreton Virus (MORV) as a Novel Oncolytic Virotherapy Platform for Liver Cancers

**DOI:** 10.1101/2022.03.10.483848

**Authors:** Bolni Marius Nagalo, Yumei Zhou, Emilien J. Loeuillard, Chelsae Dumbauld, Oumar Barro, Natalie M. Elliott, Alexander T. Baker, Mansi Arora, James M. Bogenberger, Nathalie Meurice, Joachim Petit, Pedro Luiz Serrano Uson Junior, Faaiq Aslam, Jean Christopher Chamcheu, Camila C. Simoes, Martin J. Cannon, Alexei Basnakian, Steven R. Post, Kenneth Buetow, Michael T. Barrett, Dan G. Duda, Bertram Jacobs, Richard Vile, Michael A. Barry, Lewis R. Roberts, Sumera Ilyas, Mitesh J. Borad

**Affiliations:** Department of Pathology, University of Arkansas for Medical Sciences (UAMS), Little Rock, AR, USA; Department of Microbiology and Immunology, UAMS, Little Rock, AR, USA; Department of Pharmacology, UAMS, Little Rock, AR, USA; The Winthrop P. Rockefeller Cancer Institute, UAMS, Little Rock, AR, USA; Division of Hematology and Medical Oncology, Mayo Clinic, Phoenix, AZ, USA; Department of Molecular Medicine, Mayo Clinic, Rochester, MN, USA; Division of Gastroenterology and Hepatology, Mayo Clinic, Rochester, MN, USA; Division of Infectious Diseases, Mayo Clinic Rochester, MN, USA; Steele Laboratories for Tumor Biology, Department of Radiation Oncology, Massachusetts General Hospital and Harvard Medical School, Boston, MA, USA; Center for Infectious Diseases and Vaccinology, the Biodesign Institute, Arizona State University, Tempe, AZ, USA; College of Pharmacy, University of Louisiana Monroe, Monroe, LA, USA; Computational Sciences and Informatics Program for Complex Adaptive System Arizona State University, Tempe, AZ, USA; Hospital Israelita Albert Einstein, São Paulo, Brazil

**Keywords:** cholangiocarcinoma, hepatocellular carcinoma, Morreton virus, vesicular stomatitis virus, oncolytic virotherapy

## Abstract

Morreton virus (MORV) is a novel oncolytic *Vesiculovirus*, genetically distinct from vesicular stomatitis virus (VSV). we report that MORV induced potent cytopathic effects in a panel of cholangiocarcinoma (CCA) and hepatocellular carcinoma (HCC) cell lines. In addition, high intranasal doses of MORV were not associated with significant adverse effects and were well tolerated in mice bearing liver tumor xenografts and syngeneic liver cancers. Furthermore, single intratumoral treatments with MORV (1 x 10^7^ TCID_50_) triggered a robust antitumor immune response leading to substantial tumor regression and disease control in a syngeneic CCA model, using 10-fold lower dose compared to VSV (1 x 10^8^ TCID_50_). In addition, MORV and VSV both induced prominent tumor growth delay in immunodeficient mice bearing Hep3B hepatocellular carcinoma (HCC) but not in mice bearing HuCCT-1 CCA xenografts. Our findings indicate that wild-type MORV is safe and can induce potent tumor regression in HCC and CCA animal models without adverse events via immune-mediated and immune-independent mechanisms. Further development and clinical translation of MORV as virotherapy for liver cancers are warranted.

## Introduction

Oncolytic viral vectors have exhibited potential for development as anti-cancer therapeutics.^1^ The selective replication of oncolytic viruses (OVs) inside neoplastic cells can be leveraged towards this end. OVs also manifest other multi-modal mechanisms of activity, including elucidation of host immunomodulatory effects against both viral proteins and cancer antigens.^2–8^ The genus *Vesiculovirus* in the family *Rhabdoviridae,* comprises a class of negative-sense single-stranded RNA viruses, including viruses with promising efficacy as OVs or vaccine vectors.^9,10^ Vesicular stomatitis virus (VSV), a widely studied vector, has been advanced into early phases clinical studies. ^11,12^ However, concerns for hepatic and neurological toxicities have hindered its clinical deployment for the treatment of human cancers. ^13–17^ Few other *Vesiculoviruses* vectors have been developed in recent years ^10^, most failing to achieve the level of potency of oncolytic VSV.

In this report, we describe our evaluation the oncolytic potential of Morreton virus. We conducted a first-of-its-kind evaluation of the safety profile, biodistribution and oncolytic potency of Morreton virus (MORV), a non-pathogenic *Vesiculovirus*, that is genetically distinct from VSV.^18,19^ MORV was isolated four decades ago from phlebotomine sandflies in Colombia. ^20^ Although antibodies against MORV have been detected in animals and humans in Colombia^20^, unlike VSV ^21^, MORV has not been directly associated with clinical pathology in humans or animals. Herein, we also conducted a series of experiments to assess the safety profile of MORV after intranasal administration of increasing doses of viral particles in healthy immunocompetent mice. Furthermore, we evaluated the anti-tumor efficacy of single and multiple doses of MORV in subcutaneous xenografts of cholangiocarcinoma (CCA), hepatocellular carcinoma (HCC), and an aggressive syngeneic, orthotopic mouse model of CCA. These preliminary initial findings will hopefully support further development of this promising new oncolytic vector as a novel anti-neoplastic therapy.

## RESULTS

### Ultrastructure and Genome Sequence of Morreton Virus (MORV)

A phylogenetic tree based upon MORV glycoprotein (MORV-G) compared to other known *Vesiculoviruses* shows that MORV, a non-pathogenic, bullet-shaped (length:200nm; width:75nm) negative-sense RNA *Vesiculovirus* of the family *Rhabdoviridae*, is closely related, yet genetically distinct from vesicular stomatitis virus Indiana strain (VSV-I) (Fig.1A and B; Supplementary Fig.1)^18,19^. The complete genome of MORV of 11,181 nucleotide genome of MORV is typical of *Vesiculoviruses* comprising five major genes from the 3’ to 5’ direction: nucleoprotein (MORV-N), phosphoprotein (MORV-P), matrix (MORV-M), glycoprotein (MORV-G), polymerase (MORV-L) ^18^. Similar to VSV, MORV can be grown at high titers and form relatively larger plaques in Vero cells (Fig. 2A and B).

**Figure 1.**
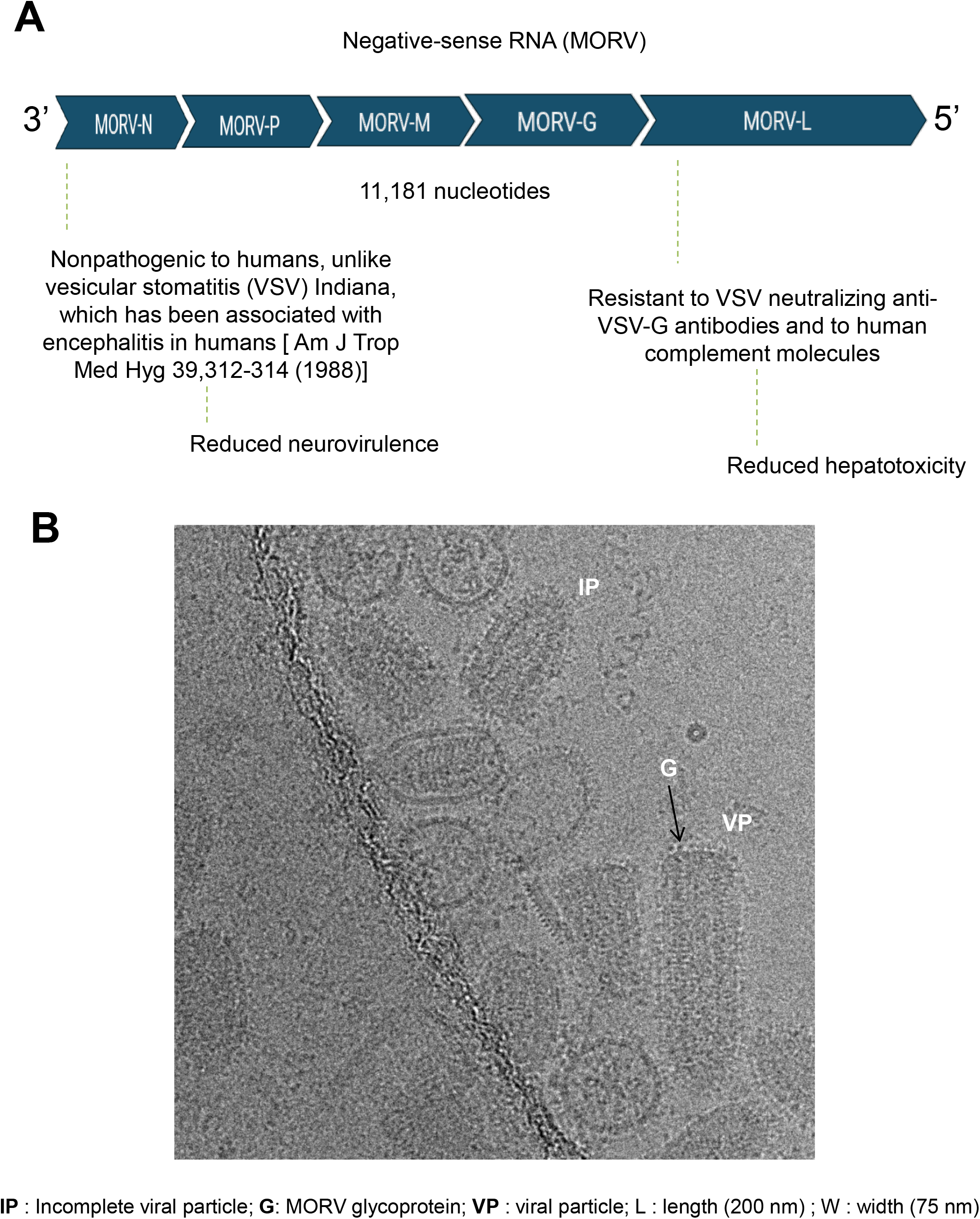
**A,** <u>Genome Organization and Ultrastructure of MORV</u>. MORV, a negative-sense RNA *Vesiculovirus* comprising five major genes (MORV-N, MORV-P, MORV-M, MORV-G, and MORV-L) and 11,181 nucleotides in genome. **B,** Transmission electron microscopy image of MORV virus, a bullet-shaped virus of 200 nm (length) and 75 nm (width). Defective interfering (DI) particles designated as incomplete particles (IP) were found after several plaque purifications. VP (viral particle) and G (glycoprotein).

**Figure 2.**
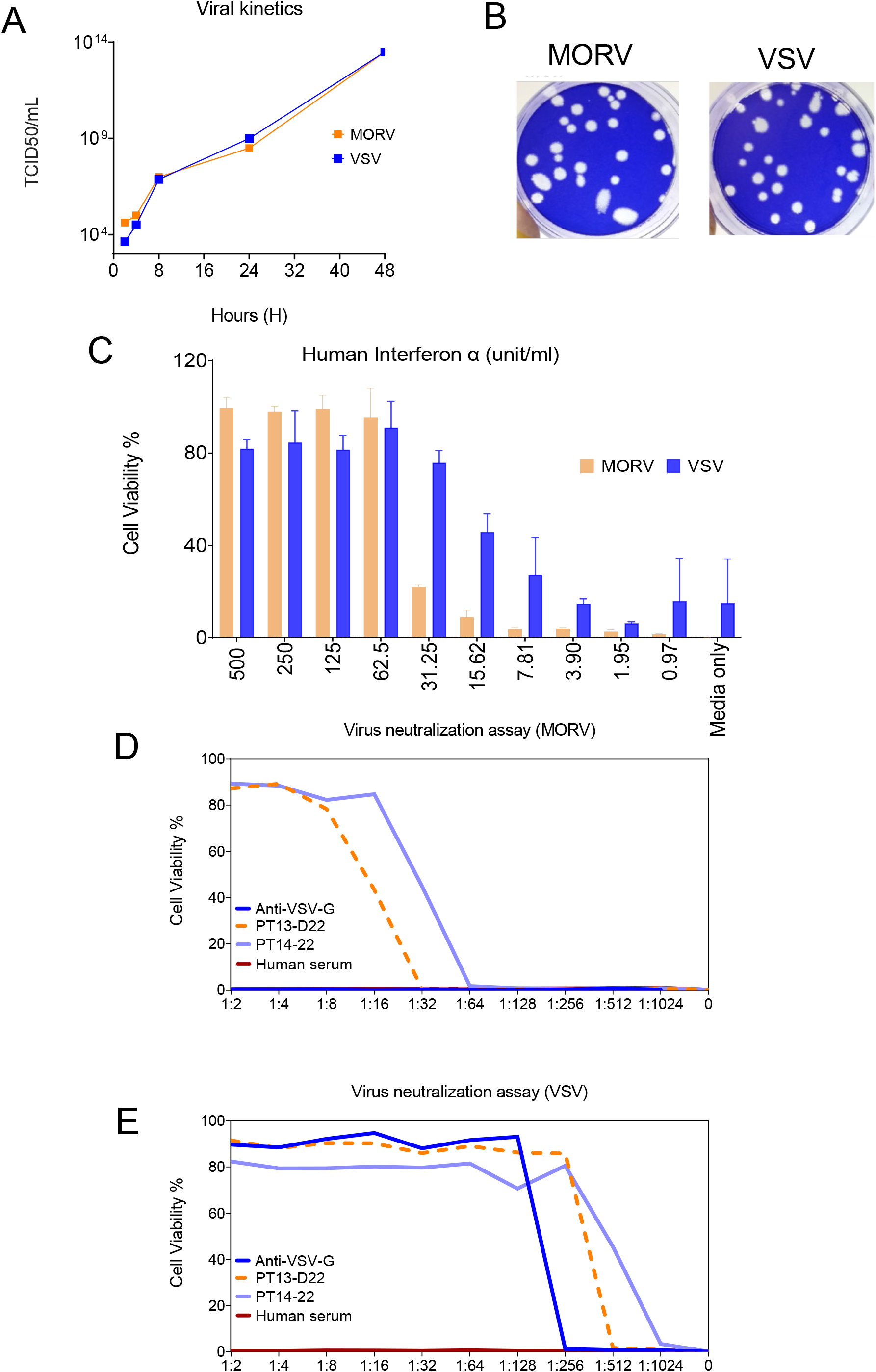
<u>Characterization of MORV</u>. **A**, Left-Viral titers of MORV and vesiculovirus stomatitis virus (VSV). Recombinant MORV and laboratory-adapted VSV were used to infect BHK-21 cells. **B**, Supernatants from infected cells were collected at different time points, and viral titer was determined using a TCID_50_ (50% tissue culture infective dose) method on Vero cells (1.5 x10^4^). Right-Serial dilutions of MORV and VSV were used to infect a monolayer of Vero cells (5 x10^5^), then stained 48 hours later with crystal violet to reveal viral plaques. **C,** A549 (2 x10^4^), interferon responsive lung cancer cells were pretreated with various concentrations of universal type I interferon-alpha (IFN-α), infected with viruses at a multiplicity of infection (MOI) of 0.01. Cell viability was measured using a colorimetric assay (MTS, Promega USA) after 48-hours post-infection. **D and E,** Normal human serum, anti-VSV glycoprotein (anti-VSV-G) antibody, serum from patients treated with VSV (PT13-D2, PT14-22) expressing human IFN-beta (VSV-hIFN-β) were evaluated for their ability to neutralize 400 TCID_50_ units of MORV and VSV in Vero cells (2 x10^4^). Data are expressed as means of triplicate from three independent experiments.

### Oncolytic MORV is Sensitive to Host Type I Interferon (IFN-α) Associated Antiviral Mechanisms

It has been established that mammalian cells respond to virus infection by synthesizing and expressing antiviral cytokines, such as type I IFN-α and IFNβ^11,12, 22–24^. Many oncolytic viruses such as VSV selectively infect and replicate preferentially in tumor cells which lack a functional IFN-associated antiviral response ^4,25–27^. Analysis of the effect of type I IFN on MORV was undertaken by treating a monolayer of A549 interferon-responsive lung cancer cells with serial concentrations of human type I IFN-α for 24 hours, followed by infection with MORV and VSV at an MOI of 5. Dose-response treatment with MORV and VSV resulted in inhibition of MORV and VSV oncolysis with differences in sensitivity to IFN-α (Fig. 2C; Supplementary Fig.2A). The lowest concentration of human IFN-α to significantly inhibit VSV-induced CPE (75% cell viability) was 31.25 units/ml. In contrast, MORV required a 2-fold higher dose of 62.5 units/ml to induce protection (85% cell viability).

### MORV is Resistant to Neutralizing Anti-VSV-G Antibodies and Normal Human Serum

A significant challenge impeding repeated systemic administration of oncolytic VSV is the rapid development of host anti-VSV neutralizing antibodies, predominantly directed against the glycoprotein (VSV-G)^28,29^. Furthermore, the proportion of patients with pre-existing immunity to VSV is not entirely known. Developing MORV as an oncolytic agent can potentially circumvent VSV immunity and allow for the development of an extended dosing approach in sequence with VSV, to target residual disease. To test this concept, anti-VSV neutralizing antibodies (Anti-VSV-G), non-immune pooled human serum, and serum from patients treated with VSV-hIFN-β (ClinicalTrials.gov: NCT01628640) were evaluated for their ability to neutralize MORV. While the anti-VSV-G antibody effectively neutralized VSV, it was ineffective against MORV-induced CPE (Fig. 2D and 2E). However, serum obtained from patients treated with VSV-hIFN-β (PT13-D22, PT14-22) neutralized MORV, but only at elevated serum concentrations (Fig.2D). Furthermore, PT13-D22 (80% cell viability at 1:80 dilution) was slightly less potent in inhibiting MORV than PT14-22 (85% cell viability at 1:160 dilution). Pooled human serum failed to neutralize MORV or VSV. Additionally, we did not find quantifiable IFN-β in serum from PT13-D22 and PT14-22, suggesting that VSV-hIFN-β did not extensively replicate in these patients. Therefore, the observed neutralization effect seen with patient serum is most likely due to polyclonal anti-VSV antibodies against viral epitopes common to VSV and MORV as previously shown^29,30^.

### MORV Induced Robust Cytopathic Effect (CPE) in Tumor Cells

Monolayers of human and murine primary liver cancer (HCC and CCA) cells, a patient-derived CCA cell line (PAX-42) and normal cholangiocytes (H69) were infected at different MOIs (1, 0.1, and 0.01), and cell viability was measured 72 hours post-infection. While, MORV or VSV did not induce a measurable cell death in normal cholangiocytes (H69), there was a cell and virus dependent variability in virus-induced CPE among tumor cell lines. Two (2) out of the 15 cell lines that were tested (HuCCT-1, PAX-42) were resistant to MORV and VSV, as manifested by 70%-100% viability at MOI of 1, 0.1, and 0.01 (Fig. 3 A, B). A similar resistance phenotype to VSV, but not MORV (35% cell viability) was also observed at MOI of 1 for EGI-1. In contrast, significant virus induced-CPE (< 10% viability at MOI of 1) was found in human SNU-1079, RBE, GBD-1, CAK-1, SNU-245, SNU-308 and SNU-869 and murine SB CCA lines, and human, HepG2, Hep3B, and Huh7, and murine RIL-175 derived clones 1 and 2 (R1LWT, R2LWT), Hepa 1-6 and HCA-1 HCC lines (Fig.3 A, B).

**Figure 3.**
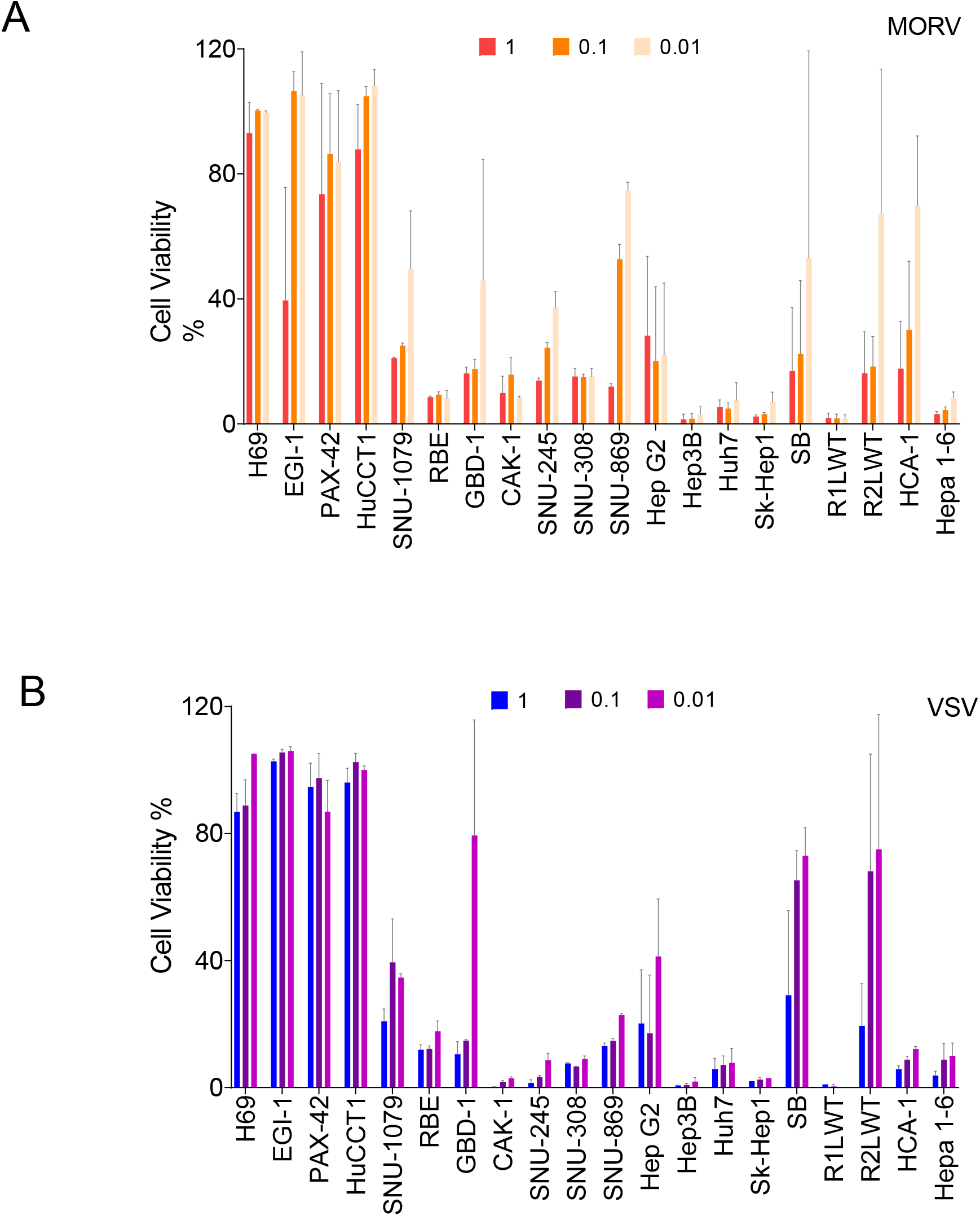
<u>Cytopathic Effect of MORV and VSV In a Panel of Liver Cancer Cells</u>. **A**, Monolayers (1.5×10^5^) of a transformed cholangiocyte cell line (H69), human CCA ( EGI-1, PAX-42, HuCCT-1, SNU-1079, RBE, GBD-1, CAK-1, SNU-245, SNU-308, SNU869), murine CCA (SB), human hepatocellular carcinoma (Hep3B, HepG2,Huh7, Sk-Hep1) and murine HCC ( R1LWT, R2LWT, HCA-1 and Hepa 1-6) cells were mock-infected or infected with MORV (**A**) and VSV (**B**) at an MOI of 0.01, 0.1 and 1. The percentage of cell viability was determined 72 hours post-infection by MTS assays (Promega, USA). The average of three independent experiments are plotted.

### The MORV-induced Cytopathic Effect in CCA is Independent of Activation and Expression of Type I IFN (IFN-β)

We showed that both MORV and VSV are highly sensitive to type I IFN. Given that MORV is closely related to VSV, we investigated whether CCA cell line permissiveness to MORV would correlate with responsiveness to type I IFN. The IFN-induction capacity of MORV and VSV were measured in supernatants of infected cells at MOI of 5 at 18 hours post-infection. While most CCA cell lines sensitive to MORV cytopathic effects failed to produce measurable IFN-β in cell culture supernatant, some expressed relatively high IFN-β (GBD-1: 600 ng/ml, SNU-308: 180 ng/ml) (Supplementary Fig.2B). Interestingly, VSV did not trigger detectable IFN-β expression in any of the CCA cells, including VSV-resistant CCA lines. This observation suggests that the resistance to CPE in some CCA may not correlate with the level of type I IFN induction. Other tumor cell line specific intrinsic mechanisms are likely to be contributory to oncolytic resistance in the absence of type I IFN secretion and signaling.

### MORV Can Infect and Lyse Low-density Lipoprotein Receptor (LDLR) Knockout Cell Line (HAP1)

The pantropic nature of VSV implies that it binds and infects host cells via attachment to a cell entry protein that is ubiquitously expressed on the surface of mammalian cells. The low-density lipoprotein receptor (LDLR) and family members have been shown to be putative primary receptors for VSV^28^. It is unknown whether other *Vesiculoviruses* such as MORV also use LDLR family members as cellular entry receptors. To address this question, we infected isogenic pairs of wild type HAP-1 (HAP1 WT) and LDLR knockout (LDLR KO) cell lines with MORV and VSV across a range of MOIs. The HAP1 cell line (Horizon Discovery, Cambridge, UK) is a human near-haploid cell line derived from the chronic myelogenous leukemia (CML) cell line KBM-7^31,32^. HAP1 LDLR KO cells have their LDLR gene disrupted by a single nucleotide excision in exon 4 using CRISPR-Cas9 technology and do not express LDLR on their surface (Fig. 4A). Surprisingly, in our evaluation both MORV and VSV efficiently infected and induced extensive cell lysis in HAP1 WT, and LDLR KO cell lines at indicated MOIs (Fig. 4B and C). This suggests that MORV and VSV cell entry are not significantly dependent on the availability of LDLR, suggesting that these viruses employ an array of receptors or receptor-independent cell entry mechanisms.

**Figure 4.**
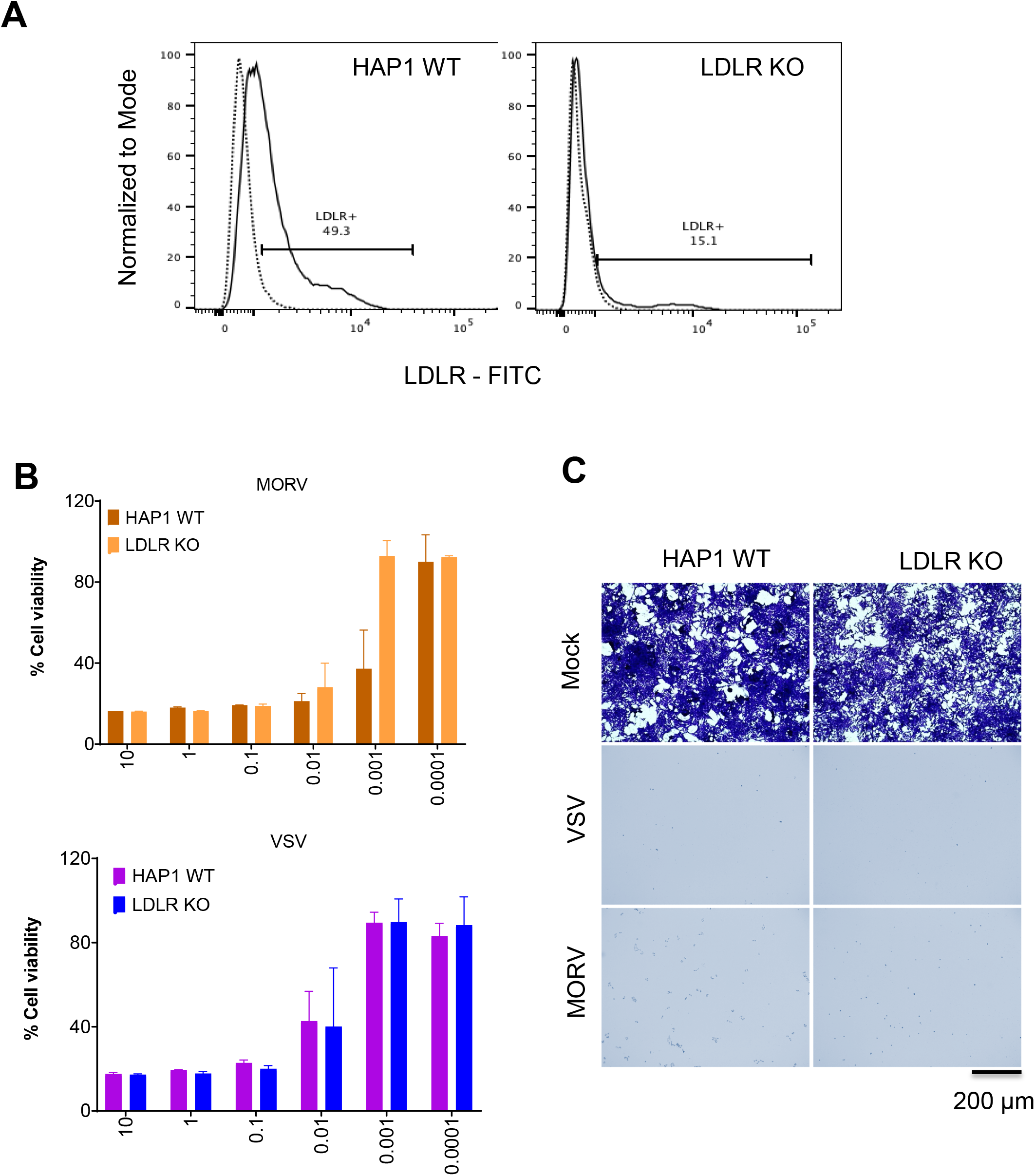
<u>Assessment of MORV and VSV Infectivity and Cytotoxicity In Low Density Lipoprotein Receptor (LDLR) Knockout Cell Line (HAP-1).</u> **A,** Expression of low-density lipoprotein receptor (LDLR) was measured by flow cytometry using fluorescein (FITC) anti-human LDLR antibody (solid line) or an isotype control antibody (dashed line). **B,** we infected HAP1 WT or LDLR KO cells (1×10^4^) cells with different MOIs (10,1, 0.1, 0.01, 0.001, and 0.0001) of MORV or VSV, and 72 hours post-infection, cell viability was measured by MTS assay. **C,** Representative images of infected or mock-infected HAP1 WT and LDLR KO cells fixed at 48 hours, stained with crystal violet (scale bar=200 µm). Data are shown as mean ± SEM from three independent experiments.

### High Dose Intranasal Administration of MORV is Not Associated With Neurotoxicity or Hepatotoxicity

In order to determine whether MORV is causally associated with brain damage and neurotoxicity in animal models, immunocompetent mice were subjected to four intranasal doses of MORV and VSV across a range of doses (1×10^7^ TCID_50_/kg, 1×10^8^ TCID_50_/kg, 1×10^9^ TCID_50_/kg, 1×10^10^ TCID_50_/kg,) or PBS. The highest dose of 1×10^10^ TCID_50_/kg corresponded to 2×10^8^ TCID_50_ total for a 20g mouse. Three mice per group were sacrificed at three days post-infection to assess short term toxicity and blood, brain, liver, and spleen were collected for further analysis. The remaining animals were monitored for 45 days by a certified veterinarian for signs of toxicity. Low to high doses of MORV and VSV were well tolerated and managed in all groups. Bodyweight decreases were not statistically significant, three days after treatment in the MORV groups, followed by recovery and a consistent increase from seven days after intranasal administration until the end of the study (Fig. 5A and B). Consistent with body weight, a thorough pathology review of brain sections of mice treated with the highest dose of viruses (1×10^10^ TCID_50_/kg) showed that MORV did not cause any overt sign of toxicity (Fig.5 C). Additionally, we performed an analysis of complete blood counts (CBC), three days after treatment. There was a decrease in the number of white blood cells, lymphocytes, granulocytes, and platelets in mice treated with MORV (Fig. 5D; Supplementary Fig. 3). In addition to virus mediated hematological changes, these changes in the blood counts may have been reflective of the hemolysis from the terminal cardiac puncture, which is known to interfere with the blood counts and other blood parameters.

**Figure 5.**
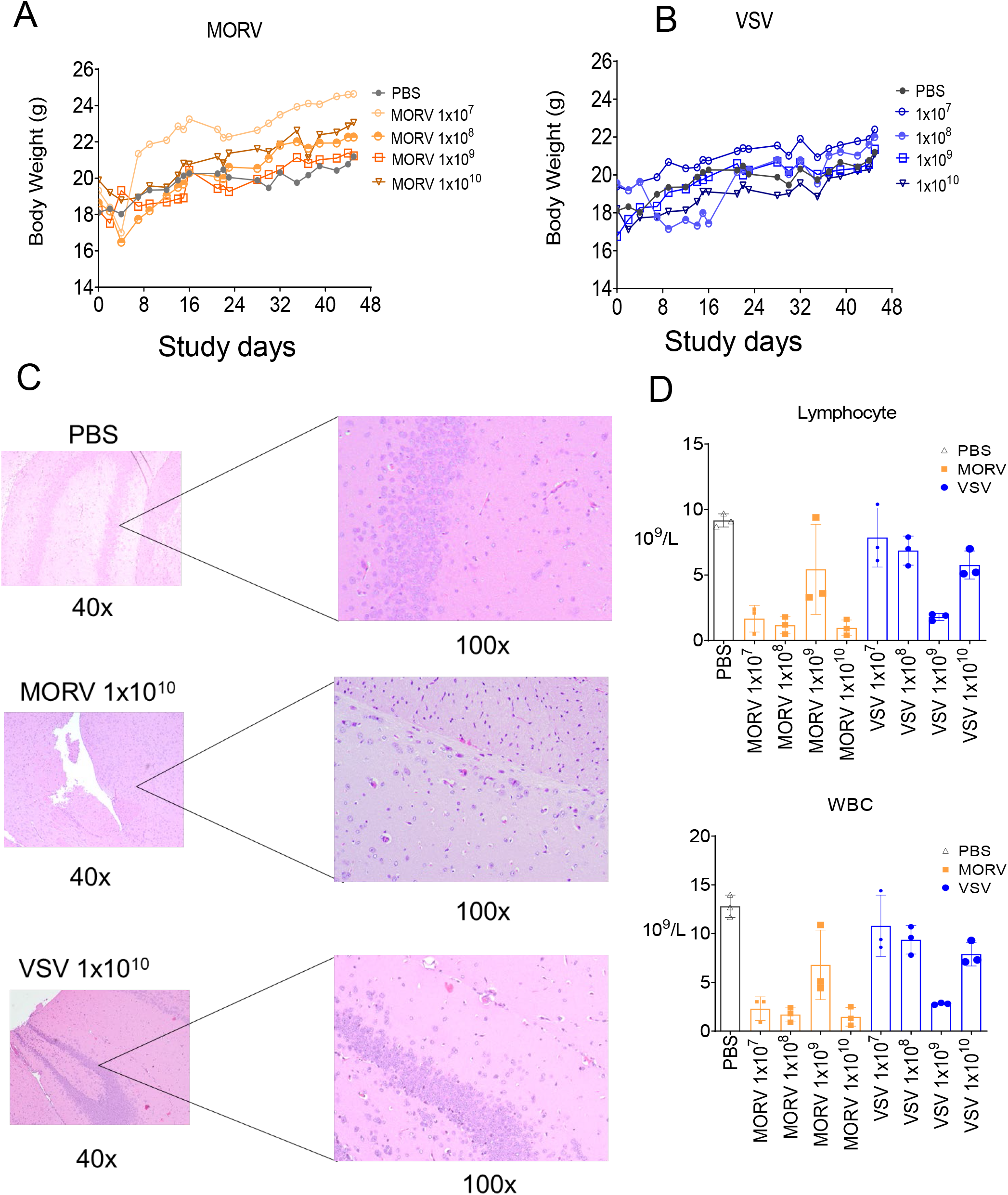
<u>Intranasal Administration of MORV and VSV in Immunocompetent Mice</u>. **A and B,** Measurement of body weight and **C**, Pictures of Hematoxylin and eosin (H&E) stained brain sections from mice treated with high dose of MORV and VSV (1×10^10^ TCID_50_/kg). **C,** complete blood count (white blood cells (WBC) and lymphocytes) values following intranasal administration of increasing doses (1×10^7^, 1×10^8^, 1×10^9^ and 1×10^10^ TCID_50_/kg) of MORV and VSV.

### MORV Did Not Extensively Replicate In Non-Tumor Bearing Immunocompetent Mice

Quantitative reverse transcription (qRT-PCR) targeting MORV-N and VSV N genes was performed on RNA samples extracted from the blood, brain, liver, and spleen. As shown in Supplementary Fig.4, MORV-N RNA was undetectable (< 10 copies per µg/Total RNA) in these tissues across the doses tested (1×10^7^-1×10^10^ TCID_50_/kg). On the contrary, we detected ∼ 30 copies/Total RNA of VSV-N RNA in the brain at 1×10^9^ TCID_50_/kg and 1×10^10^ TCID_50_/kg as shown in Supplementary Fig.4. Collectively, these data indicate that intranasal administration of high doses of MORV (up to 1×10^10^ TCID_50_/kg) did not elicit significant toxicity (100% survival after 45 days) in the treated animals, or result in physical impairment (Supplementary Table 3).

### Heterogeneity In Anti-tumor Efficacy of MORV In Xenograft Models of Human Liver Cancer

As delineated above, the level of virus-specific receptor expression, type I IFN production, or viral progeny production *in vitro* by infected cancer cell lines do not fully explain variable susceptibility to MORV-induced cell death. Therefore, to assess whether a tumor cell line that is resistant to MORV-induced CPE *in vitro* would display the same phenotype *in vivo*, resistant (HuCCT-1, cholangiocarcinoma) and sensitive (Hep3B, hepatocellular cancer) cancer cell lines were subcutaneously implanted into female athymic nude (NU/J) mice (n=7) (Jackson Laboratories, USA). After tumors reached a treatable size, single doses or weekly intra-tumoral injections of MORV and VSV at 1 x 10^7^ TCID_50_ units were performed. Body weight, tumor volume, and weight, and clinical parameters were recorded three times a week. In the human HuCCT-1 CCA model, MORV (*p*= 0.32) and VSV (*p*= 0.26) failed to induce significant reduction of tumor growth (Fig. 6A and B). In contrast, a single injection with MORV (*p*= 0.0068) and VSV (*p*= 0.016) in human Hep3B HCC xenografts was sufficient to induce significant tumor growth delay (∼80%), (Fig. 6C and D). There were no adverse events, or virus-related toxicity noted in the study. This implies that MORV can infect and lyse tumor HCC cells *in vivo*, but not CCA line *in vivo*. Moreover, a dense stroma characterizes CCAs^33^, which could also have reduced virus intra-tumoral penetration and bioavailability.

**Figure 6.**
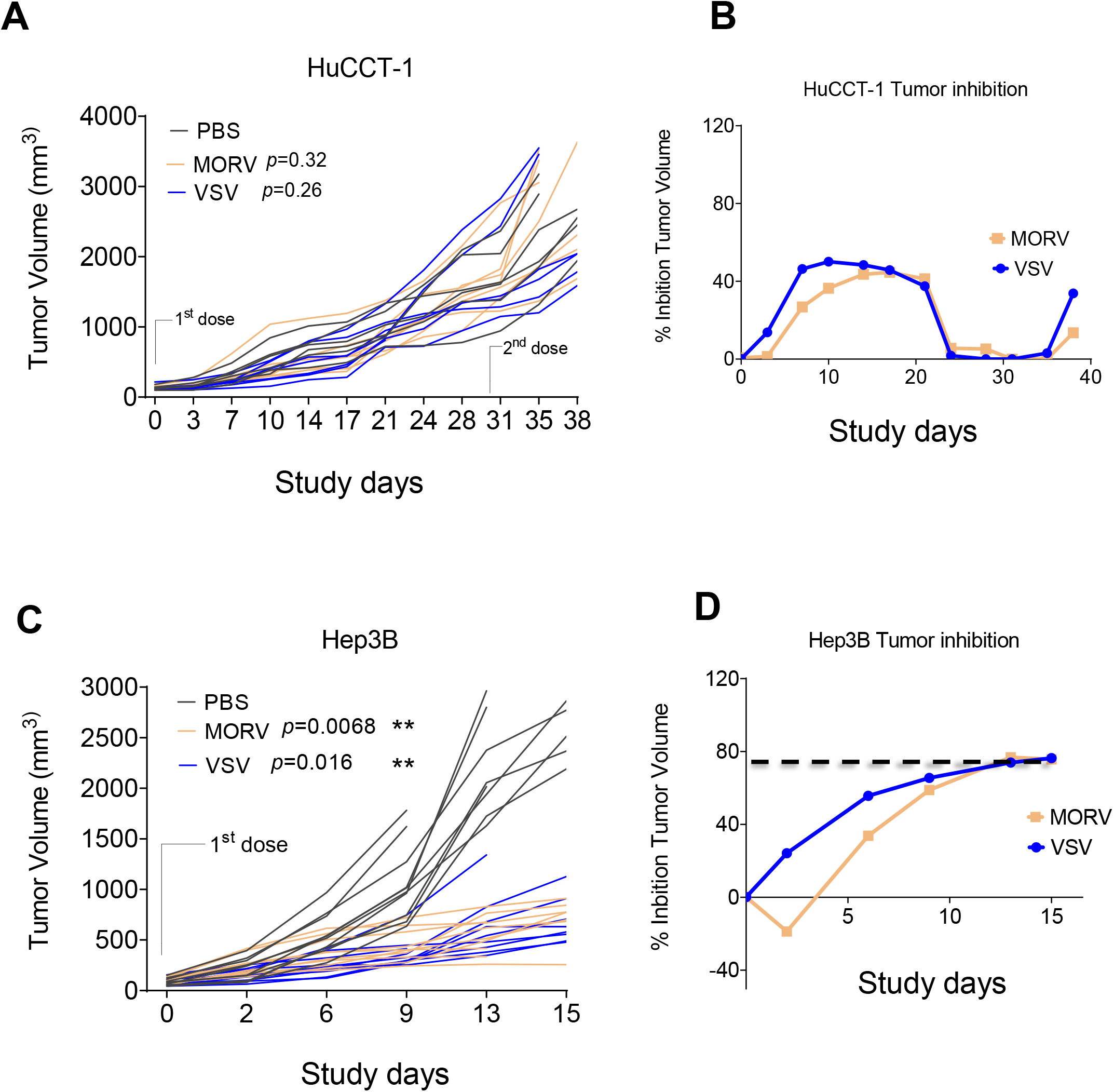
*<u>In Vivo</u>* <u>Anti-tumor Efficacy of MORV and VSV in Xenograft Mouse Models of Cholangiocarcinoma (CCA) and Hepatocellular Carcinoma (HCC).</u> Single or multiple doses (2) of 1 × 10^7^ TCID_50_ of MORV or VSV were injected intratumorally into HuCCT-1(CCA) and Hep3B (HCC) tumor-bearing mice. Analysis of the effect of MORV and VSV on tumor growth and tumor inhibition in HuCCT-1 (**A**), and Hep3B (**B**) xenografts.

### Low Dose of MORV Induces a Significant Reduction of Tumor Burden Compared to VSV In an Aggressive Syngeneic, Orthotopic Mouse Model of CCA

To investigate the anti-tumor effect of MORV, murine bile duct cancer cells (SB) derived from an oncogene-driven murine model of CCA were surgically implanted into livers of immune competent mice as previously described.^34,35^ The SB orthotopic model is Yes-associated protein (YAP)-driven with considerable overlap at the messenger RNA level with human intrahepatic CCA, and as such is a representative model for phenotype and progression recapitulation of human CCA ^33,36,37^. Before tumor cell implantation, we confirmed that MORV can efficiently infect and kill SB cells *in vitro* using real-time analysis of viral-induced apoptosis (Supplementary Fig. 6A). Fourteen days after orthotopic implantation of SB cells, single intraperitoneal (IP) doses of PBS, 1×10^7^ TCID_50_ or 1×10^8^ TCID_50_ units of MORV, or VSV were administrated to mice (Fig. 7A). Four weeks following SB cell implantation, mice were sacrificed, and cardiac blood and tumors were collected for downstream analysis. A significant reduction of malignant nodule size was observed in the MORV-treated mice at 1×10^7^ TCID_50_ (*p*=0.0001), which was 10-fold lower than the dose of VSV (1 x 10^8^ TCID_50_) showing similar tumor regression (*p*=0.0018) and disease control (Fig. 7B). Elevated serum alanine aminotransferase (ALT) and alkaline phosphatase (ALP) levels are hallmark of liver and biliary tract damage. There was no significant difference in the ALT and ALP levels with MORV and VSV compared to vehicle (Fig. 7 D and E). Additionally, the adjacent mouse liver showed no TUNEL staining indicating, an absence of liver toxicity (Supplementary Fig. 6B). No significant changes were observed in serum IFN-α and IFN-β, serum albumin, cholesterol, blood urea, and expression levels of MORV-N and VSV-N genes in tumor nodules (Supplementary Fig. 7B and C).

**Figure 7.**
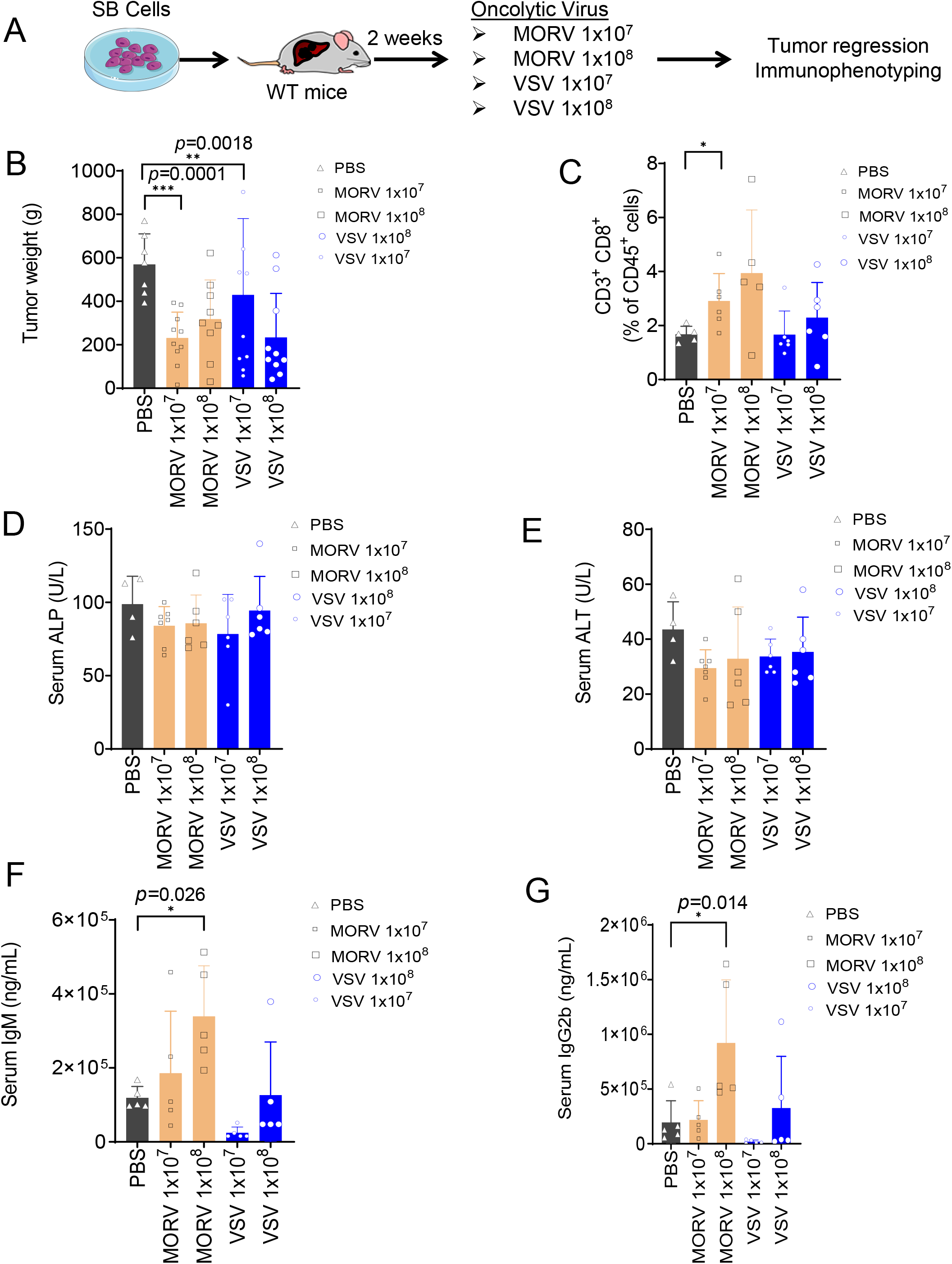
<u>MORV Reduced Tumor Burden At a Lower Dose Compared to VSV In Murine CCA.</u> **A,** Overview of the *in vivo* experiment**. B,** Tumor weight of mice treated with phosphate-buffered saline (PBS), MORV, and VSV at 1 x10^7^ TCID_50_ or 1×10^8^ TCID_50_. **C,** Frequency of tumor-infiltrating CD3+ CD8+ CTLs. **D, E, F and G, graphs** showing changes in serum ALP, ALT, IgM and IgG2b.

### MORV Antitumor Efficacy is Associated With a Significant Increase in Intra-Tumoral CD8^+^ Cytotoxic T Lymphocytes (CTLs)

Oncolytic vesiculoviruses exert their anti-tumor actions through induction of direct cytotoxicity in tumor cells and modulation of immune-mediated mechanisms. Here, we investigated MORV treatment effects in an orthotopic syngeneic model of intrahepatic CCA by assessing both tumor-infiltrating immune cells (lymphocytes, macrophages, and myeloid-derived cells) and antibody response to virus infection using multi-color flow cytometry and mouse immunoglobulin isotyping multiplex assay. The immune response to IP administration of MORV was significantly more potent than that observed in the VSV-treated group. We observed a significant increase in CD3^+^ CD8^+^CTLs (*p*=0.029), IgG2B (*p*=0.014), IgG3 (*p*=0.0084), and IgM (*p*= 0.0256) in the MORV treated groups compared to VSV, which correlates with the robustness of the antiviral and anti-tumor immune responses resulting in significant reduction of tumor weight and disease control (Fig.7C, F and G, Supplementary Fig. 9).^30,38^ M2-like tumor-associated macrophages (TAMs) and myeloid cells are known to exert their immunosuppressive actions via expression and interaction of PD-L1 with PD-1 on the surface of CD8+ T cells.^39,40^ Furthermore, the increased TAM frequency in stroma-rich malignancies such as CCA is well documented.^33^ We show that infection with MORV did not increase the frequency of TAMs (M2-like) and monocytic myeloid-derived suppressor cells (M-MDSCs) or granulocytic (G)-MDSCs (Supplementary Fig. 8). These results suggest that systemic administration of MORV is associated with an effective and robust anti-tumor immune response mainly via increasing CD8^+^ T cells. In addition, treatment with MORV increased the number of TUNEL-positive apoptotic tumor cells in both MORV and VSV treated groups compared to the control group with no overt effects in adjacent normal liver cells (Supplementary Fig.6B). We did not detect the MORV-N gene in tumor nodules at the end of the study, suggesting that transient intra-tumoral replication of MORV can induce and sustain durable anti-tumor immunity.

## DISCUSSION

Vesicular stomatitis virus (VSV) has been under development as an oncolytic agent and vaccine vector ^11,29,41,42^. However, clinical translation of VSV has been hindered by several factors, including need for use of attenuated vectors such as VSV-human interferon beta, that would mitigate concerns raised by observations of neurovirulence ^16,17,21, 43–45^, and liver toxicity^41^ in laboratory animals. Furthermore, there have been also reported cases of VSV-induced encephalitis in humans^21^. In this study, we conducted a thorough evaluation of the toxicity, biodistribution and the anti-tumor activity of Morreton virus (MORV), a novel oncolytic *Vesiculovirus,* genetically distinct from vesicular stomatitis virus (VSV), in liver cancer cell lines (HCC and CCA), tumor xenografts, and an immunocompetent syngeneic CCA model.

Here, we demonstrated that intranasal administration of MORV did not cause significant neurological adverse events in mice, including encephalitis. Moreover, MORV potently induced CCA and HCC cell death *in vitro*, in subcutaneous tumor xenografts and in the syngeneic orthotopic model of CCA. MORV was highly sensitive to type I interferon (IFN) bioactivity,^46^ yet exhibited a robust lytic cycle in cancer cells and produced high viral titers in culture. These observations suggest that the tumor selectivity nature of MORV is due to inability of some tumor cells to activate a functional host IFN-associated antiviral response as seen for other oncolytic *Vesiculoviruses*. ^2–8^ However, we also observed that there was not a clear association between expression of cellular type I IFN and permissiveness to MORV in CCA cell lines. Many groups have studied IFN response induced by vesiculoviruses.^47^ How these viruses trigger activation of the type I IFN signaling pathway may not be shared within the family, but could be rather a virus-specific mechanism that needs to be further investigated.

MORV induced-cytopathic effect (CPE) in cancer cells was not affected by exposure to anti-VSV glycoprotein (anti-VSV-G) antibody or normal human serum. Conversely, serum from liver cancer patients treated with VSV-hIFNβ, a VSV-based oncolytic vector currently being evaluated in a Phase I clinical trial (ClinicalTrials.gov: NCT01628640), had modest inhibitory activity against MORV infectivity (concentrations ≥ 1:80), but strongly neutralized VSV infection (concentrations ≥ 1:2560). These results suggest that MORV may share similar antigenic determinants with VSV to which antibodies may specifically bind and partially neutralize MORV activity.

Although the cellular entry receptor for MORV is unknown, it is possible that receptors are shared among the *Vesiculovirus* family. VSV is known to use the LDLR and family members. Intriguingly, both MORV and VSV were able to infect and lyse HAP-1 LDLR KO cells efficiently. This result shows that permissiveness to VSV is not affected by the availability of LDLR and that MORV may also use a different cellular entry protein. Furthermore, this suggests that other cellular receptors or entry mechanisms are implicated in VSV and MORV susceptibility.

Overall, infection with MORV triggered a significant (30%-90%) CPE in HCC and CCA cells *in vitro*, and MORV induced CPE was comparable to that of VSV. Several human and the mouse CCA tumor cells were permissive to MORV whereas normal cholangiocytes (H69 cells) were not.

High intranasal doses of MORV in healthy immunocompetent mice were found to be safe and tolerable. The short-term (three days post-infection) toxicity analyses showed a slight decrease in lymphocyte count, granulocytes, and platelets in mice treated with MORV. Previous studies have shown that infection with vesiculoviruses, especially VSV is associated with interferon-mediated lymphopenia in laboratory mice ^48,49^. In this study, no adverse clinical signs, including hepatotoxicity, neurotoxicity, or significant loss of body weight were noticed.

Most oncolytic viruses have shown heterogeneous efficacy across multiple cancer types. Furthermore, extended dosing strategies have been shown to effectively target and eliminate residual disease and enhance oncolytic effects.^50^ Therefore, we evaluated the antitumor efficacy of a single and multiple intra-tumoral doses of MORV and VSV in liver tumor xenografts. The anti-tumor efficacy in the CCA and HCC xenografts correlated with the *in vitro* CPE observed in HuCCT-1(resistant) and Hep3B (sensitive) lines. Specifically, while two doses of MORV and VSV treatments were ineffective against the HuCCT-1 tumor xenografts, MORV induced substantial tumor regression in the Hep3B xenograft models.

Surprisingly, a single dose resulted in a robust antitumor immune response, characterized by a significant increase in tumor-infiltrating CD8^+^ cytotoxic CTLs leading to prominent tumor regression in tumor weight in the syngeneic orthotopic SB mouse model of CCA. Moreover, MORV exerted superior antitumor activity at a 10-fold lower dose than VSV. These results strongly suggest that MORV can stimulate and sustain an antitumor immune response at a relatively lower dose than VSV, thus reducing concerns related to toxicities associated with high doses of oncolytic viruses.

Together, these data demonstrate that robust efficacy and safety can be achieved using MORV in animal models. We have successfully shown that MORV is sensitive to host type I IFN, resistant to anti-VSV-G antibody, and has robust cell-killing activity against CCA cells *in vitro* and animal models. We have also shown that the heterogeneity in the cytotoxicity effect seen with MORV *in vitro* correlates with its *in vivo* efficacy in CCA xenografts. As demonstrated by others and in this study, the oncolytic activity of OVs is predominantly due to their ability to elicit an active immune response against viral antigens and tumor-specific antigens ^51^. Subcutaneous human tumor xenografts are well-established models that faithfully reflect the clinical outcomes in human disease ^52^; however, these models lack a functional immune system; thus, significantly limit our ability to fully evaluate the immunomodulatory potential of OVs. In addition, the dense stromal component of the CCA tumor microenvironment ^33^ also contributes to altering the efficient intra-tumoral biodistribution of viral particles.

Therefore, further research is warranted to identify biomarkers that could predict the likelihood patient to benefit from MORV oncolytic virotherapy. Nonetheless, these preliminary results provide the rationale for future studies to fully characterize MORV as a potent novel oncolytic agent to treat human bile duct cancers. These studies also lay the groundwork for MORV ultimately being evaluated in first-in-human studies.

## MATERIALS AND METHODS

### Cell Lines

This study used a panel of 9 human cholangiocarcinoma (CCA) cell lines (HuCCT-1, EGI-1, CAK-1, SNU-245, SNU-308, SNU-869, SNU-1070, RBE, GBD-1), 1 carcinoma of the ampulla of Vater, a transformed cholangiocyte cell line (H69), a murine CCA (SB), 4 human hepatocellular carcinoma (HCC) cell lines (HepG2, Hep3B, Huh7, Sk-Hep-1) and 3 murine HCC cell lines (R1LWT, R2LWT, HCA-1). All cell lines were cultured at 37° C with 5% CO_2_ in media supplemented with L-glutamine and antibiotic agent (100 µg ml^-1^ penicillin and 100 µg ml^-1^ streptomycin). EGI-1, CAK-1, SNU-245, SNU-308, SNU-869, SNU-1070, RBE, GBD-1 were maintained in RPMI 1640 medium with 10% fetal bovine serum (FBS). PAX-42, HuCCT-1, H69, BHK-21 (Baby Hamster kidney fibroblast), A549 (human lung tumor cells), and Vero (African green monkey kidney) cells were maintained in Dulbecco’s Modified Eagle’s Medium (DMEM) with 10% fetal bovine serum (FBS). CAK-1, GBD-1, H69 were gifts from Dr. Gregory Gores at Mayo Clinic, Rochester, MN. A patient-derived cell line (PAX-42, grown in DMEM with 10% FBS and 100 ug ml^-1^ Primocin) was a gift from Dr. Mark Truty at Mayo Clinic, Rochester, MN. HAP1 parental cell line and LDLR-KO cell line were obtained from Horizon and maintained in Iscove’s Modified Dulbecco’s Medium (IMDM) supplemented with 10% FBS, 1 % antibiotic. We obtained the HuCCT-1 cell line from the Japanese Collection of Research Bioresources (JCRB). EGI-1 was purchased from the German Collection of Microorganisms and Cell Cultures (DSMZ). SNU-245, SNU-308, SNU-869, SNU-1079 were obtained from the Korean cell line bank (KCLB). RBE was purchased from the National Bio-Resource Project of the MEXT, Japan (RIKEN). BHK-21, A549, and Vero cells were obtained from the American Type Culture Collection (Manassas, VA). Hep3B, HepG2, Huh7, Sk-Hep1 and Hepa 1-6 were purchased from the American Type Culture Collection (ATCC, Manassas, VA) and were grown in DMEM with 10% FBS and RPMI with 10% FBS, respectively. The two clones derived from the RIL-175 cell line (R1LWT, R2LWT), HCA-1 and SB cells were grown in DMEM with 10% FBS.

### Oncolytic Viruses

We obtained Morreton virus (MORV) from the University of Texas Medical Branch (UTMB) World Reference Center for Emerging Viruses and Arboviruses (WRCEVA). A laboratory-adapted viral clone of MORV was generated using sequential plaque purifications on Vero cells (ATCC, Manassas, VA). RNA-sequencing was applied to confirm the full-length MORV genome (Iowa State University Veterinary Laboratory (ISU VDL)). The full-length MORV genome (11,181 nucleotides) comprising genes encoding for the nucleoprotein (MORV-N), phosphoprotein (MORV-P), matrix protein (MORV-M), glycoprotein (MORV-G), and RNA-directed RNA polymerase L protein (MORV-L), was synthesized (Genscript, USA) from the laboratory-adapted viral clone of MORV and subcloned into plasmid (pMORV-XN2). pMORV-XN2, along with helper plasmids (pMORV-P, pMORV-N, and pMORV-L) as previously described ^50^. These plasmids were used to express the antigenomic-sense RNA of MORV under the bacteriophage T7 promoter to generate recombinant MORV (rMORV). However, in this study we used wild-type MORV and not recombinant rMORV. Vesicular stomatitis virus (VSV) was rescued on BHK-21 cells from a plasmid (pVSV-XN2) containing the VSV Indiana genome serotype. All viruses were rescued using a vaccinia rescue system and propagated and titrated on BHK-21 cells as previously described^53,54^. Sucrose density gradient centrifugation was used to obtain purified viral particles (VSV, MORV and recombinant MORV) before *in vitro* and *in vivo* studies.

### Evolutionary Analysis of MORV Glycoprotein (MORV-G) Amino Acids Sequence By Maximum Likelihood Method

All viral glycoproteins sequences were retrieved from www.uniprot.org. We inferred the evolutionary history by using the Maximum Likelihood method and JTT matrix-based model ^55,56^. The tree with the highest log likelihood is shown. The percentage of trees in which the associated taxa clustered together is shown next to the branches. Initial tree(s) for the heuristic search were obtained automatically by applying Neighbor-Join and BioNJ algorithms to a matrix of pairwise distances estimated using the JTT model and then selecting the topology with superior log likelihood value. The tree is drawn to scale, with branch lengths measured in the number of substitutions per site. We conducted an evolutionary analysis in MEGA X 5 using 15 amino acid sequences comprising 574 positions in the final dataset ^57^.

### Transmission Electron Microscopy (TEM)

MORV was propagated in BHK-21 cells and the supernatants harvested approximately 48hrs post infection and clarified by centrifugation at 1,500g in a benchtop centrifuge for 20mins. 2ml of 20% (w/v) sucrose was layered on top of 2ml of 60% (w/v) sucrose in an ultraclear centrifuge tube compatible with the SW28 rotor. MORV containing supernatants were layered on top of this 1-step sucrose gradient and centrifuged at 100,000g using an SW28 rotor for 2 hours. The visible virus band was harvested from between the sucrose layers by pipetting from the meniscus. MORV was then placed in a centrifuge tube compatible with an SW41 Ti rotor and diluted in 20mM Tris, 150mM NaCl pH7.8 buffer to fill the tube and centrifuged at 100,000g for 1.5hrs. The supernatant was discarded and the virus pellet was resuspended in 100ul of 20mM Tris, 150mM NaCl pH7.8 buffer. Two ul of this sample was applied to R2/2 UltrAUfoil grids and blotted manually before plunge freezing in liquid ethane. The vitrified specimen was imaged using a FEI Titan Krios transmission electron microscope (TEM, ThermoFisher) operating at an accelerating voltage of 300 keV with a Gatan K2 Summit direct electron detector (DED) camera (Pleasanton, CA).

### Viral Plaque Formation Assays

Serial dilutions (1:100 to 1:10^8^) of MORV and VSV were used to infect Vero cells (5 x 10^5^) in tissue culture dishes (60 mm) for one-hour. Cells were then, washed with phosphate-buffered saline (PBS), and overlaid with 1.5% bacteriological agar (Sigma, USA). Forty-eight (48h) hours post-infection, infected cells were stained with crystal violet, and plaques were counted to determine the plaque-forming units per ml (PFU/ml) B) Visualization of MORV and VSV plaques (1:10^5^ dilution) following crystal violet staining was performed using an Invitrogen EVOS FL Auto Imaging System.

### Interferon Sensitivity Assays

Interferon-sensitive lung cancer cells (A549) were seeded in a 96-well plate at a density of 2×10^4^ cells/well and cultured overnight. Twenty-four hours post-infection, cells were pretreated with different concentrations of Universal type I IFN-α was added directly into the culture medium. After overnight incubation, fresh medium containing Universal Type I IFN-α (Catalog No. 11105-1; PBL Assay Science, USA) was added, and cells were infected with MORV or VSV at an MOI of 0.01. Cell viability was assessed using a Cell Titer 96 AQueous One Solution Cell Proliferation Assay (Promega Corp, Madison, Wisconsin, USA). Absorbance measurements at 490 nm were normalized to the maximum read per cell line, representing 100% viability. Data are shown from three independent experiments. For all cell viability experiments, absorbance was read using a Cytation 3 Plate Reader (BioTeK, Winooski, VT, USA). Data are expressed as means of triplicates from three independent experiments +/- SEM.

### Type Interferon (IFN-β) Production Assays

Twenty thousand (2 x 10^4^) cells were plated per well in 96-well plates and infected with MORV or VSV at MOI of 5. Cell supernatants were harvested at 18 hours post-infection, and IFN-β levels were measured using a VeriKine-HS Human IFN Beta TCM ELISA Kit (Catalog No. 41435-1; PBL Assay Science, USA) as recommended by the manufacturer.

### Virus Neutralization Assays

Four hundred (400) TCID_50_ of MORV or VSV were incubated with increasing dilutions (1:10, 1:20, 1:40; the final antibody dilution was 1:10,240) of a specific polyclonal rabbit antibody against VSV-G, VSV-N and VSV-M (Imanis, USA), serum from patients (PT13-D22, PT14-22) treated with VSV-interferon beta in the context of a clinical study (ClinicalTrials.gov: NCT01628640) and normal human serum in 96-well plates and incubated one hour at 37 degree Celsius with 5% CO_2_, followed with an addition of 2 x 10^4^ Vero cells directly into the wells of the plate containing virus plus antibody. Cell viability was measured 24 hours post-incubation using an MTS assay (Promega, Madison, WI, USA) as indicated above. Data are expressed as means of triplicates from three independent experiments +/- SEM.

### Cell Viability Assays

For all cytotoxicity assays (96-well format), 1.5×10^4^ cells were infected with MORV and VSV at the indicated multiplicity of infection (MOI) of 1, 0.1, and 0.01 in serum-free Gibco Minimum Essential Media (Opti-MEM). Cell viability was determined using Cell Titer 96 AQueous One Solution Cell Proliferation Assay (Promega Corp, Madison, Wisconsin, USA). Data was generated as means of six replicates from two independent experiments +/- SEM.

### Visualization of Virus-induced Cytopathic Effects in Cholangiocarcinoma Cells

MORV and VSV were used to infect 5 × 10^5^ adherent cells per well in 6-well plates at an MOI of 0.1. Cells were incubated at 37°C until analysis. At 72 hours after infection, cells were fixed with 5% glutaraldehyde and stained with 0.1% crystal violet to visualize cellular morphology and remaining adherence indicative of cell viability. Pictures of representative areas were taken.

### Real-time Analysis of Virally Induced Apoptosis

Murine cholangiocarcinoma (SB) cells (1×10^4^ per well) were plated in 96-well plates and rested overnight. The next day, serial dilutions of virus starting at MOI of 10 were added to wells in triplicate. Annexin V Red (Essen Bioscience) was added to each well, including control wells without virus. Plates were imaged every 6 hours in the IncuCyte S3 system (Essen Bioscience, USA), recording both phase and red fluorescence images for a total of 48 hours. Total red area for each well was quantified and normalized for the entire experiment. Control wells were used to subtract background and as representative of 100% viability. Data was plotted as mean ± SEM. Representative images are included from the 48 hours timepoint with either virus-treated (MOI = 10) or control wells (mock-infected).

### Flow Cytometry

For flow cytometry, 1 x 10^6^ cells of HAP1 parental (WT) and LDLR KO were resuspended in phosphate-buffered saline (PBS) and plated in 96-well format and washed twice at 350g x 3 min. This step was followed by incubation with 3% bovine serum albumin (BSA) for 30 min at RT to block non-specific staining. Cells were washed and primary antibody (LDLR antibody, Proteintech 10785-1-AP, USA) was added at 0.2 ug/test. Primary antibody was incubated for 2 hours at room temperature then cells were washed twice with 2% FBS in PBS. Secondary antibody (Goat anti-rabbit IgG-FITC, Santa Cruz #sc-2012, USA) was added at 1:400 and incubated for one hour at room temperature (RT). Cells were washed twice with 2% FBS in PBS and resuspended in 2% formalin before being acquired on an LSRFortessa (BD). Data was collected in the FITC and scatter channels. Data were analyzed in FlowJo version 10.7.1 (BD). Isotype control samples were used to set positive gates (at 5% on isotype samples) and applied to full-stained samples.

### Susceptibility and Effect of MORV and VSV In a Low-density Lipoprotein Receptor (LDLR) Knockout Cell Line (HAP-1)

HAP1 cell line (Horizon Discovery, Cambridge, UK) is a near-haploid human cell line derived from chronic myelogenous leukemia (CML) cell line KBM-7. HAP1 cells had a single base pair deletion in exon 4 of their low-density lipoprotein receptor (LDLR) gene using CRISPR-Cas9 technology. Ten thousand (1×10^4^) cells of HAP1 WT or LDLR KO cells per well were plated in 96-well plates and rested overnight. The following day, cells were infected with serial dilutions of the indicated virus, starting with a MOI of 10. After 72 hours, cell viability was measured by MTS assay (Promega, USA) as described above. Data are shown as mean ± SEM from three independent experiments.

### ANIMAL STUDIES

Under a Mayo Clinic Institutional Animal Care and Use Committees (IACUC) approved protocol, we conducted *in vivo* evaluations described below.

### Toxicology and Biodistribution of Virus Administrated Via the Nasal Route

To determine whether treatment with MORV and VSV could be associated with neurotoxicity, 78 female C57BL/6J mice (n=6 mice per group), including control, were intranasally administrated with 10 microliters in each nostril of phosphate-buffered saline (PBS) or low to high doses (1×10^7^, 1×10^8^, 1×10^9^ and 1×10^10^ TCID_50_/kg) of MORV or VSV. Body weight, temperature, behavior, and clinical signs were monitored by a board-certified veterinarian at least three times a week for to detection of any signs of toxicity. Three days post-infection, three mice per group were sacrificed, and tissues were collected (brain, liver, and spleen) for short-term toxicity evaluation and viral biodistribution. The remaining mice were monitored for forty-five (45) days. Mice body weights and clinical observations were recorded at least three times per week for the study duration.

### Blood Tests

Blood was collected from the submandibular vein (cheek bleed) and cardiac puncture at days 3 and 45, respectively. Blood was collected in BD Microtainer tubes with ethylenediaminetetraacetic acid or lithium heparin (Becton, Dickinson and Company, Franklin Lakes, New Jersey, USA) for complete blood counts or in BD Microtainer SST tubes (Becton, Dickinson, and Company) for serum analysis. CBC analysis was performed in an Abaxis Piccolo Xpress chemistry analyzer (Abaxis, Union City, California, USA), and blood chemistry analysis was done in a VetScan HM5 Hematology Analyzer (Abaxis).

### Viral RNA Extraction and Quantitative RT-PCR

At 72 hours post-infection, 39 female C57BL/6J mice (n=3 mice per group) from PBS-treated, MORV, and VSV infected were sacrificed. RNA was extracted from tissues and blood using RNeasy® Plus Universal Mini Kit (Cat. 73404, Qiagen, USA) and QIAamp® Viral RNA (Cat. 52904, Qiagen, USA). A One-step multiplex quantitative reverse transcription (qRT-PCR) targeting MORV N and VSV N genes (IDT, USA) was used to quantify virus gene copies number in the brain, liver, spleen, and blood using a Light Cycler ® 480 RNA Master Hydrolysis (Roche Diagnostics, USA) as recommended by the manufacturer. All qRT-PCR assays, including detection of viral genes in the tumor (SB *in vivo* model) were conducted on a Light Cycler ® 480 II with the following primers and probes: VSV-N Forward primer: 5’- TGATAGTACCGGAGGATTGACGAC-3’; VSV-N Reverse primer: 5’- CTGCATCATATCAGGAGTCGGT-3’; VSV-N(Probe): 5’-FAM- TCGACCACATCTCTGCCTTGTGGCGGTGCA-ZEN/IBFQ-3’; MORV-N Forward primer: 5’-CCCCAATGCAGGGGGACTCACAAC-3’; MORV-N Reverse primer: 5’-TAGCAACATGTCTGGGGTGGGC-3’; MORV-N (Probe): 5’- HEX-TCTACAACATCTCGTCCTTGAGGAGGAGCA-ZEN/IBFQ-3’; MORV-G Forward primer: 5’-TCGCAGGACCCATCATTCCTC-3’; MORV-G Reverse primer: 5’-CAACATCTTCGTAGGGGTAC-3’; MORV-G (Probe): 5’-Cy5- CGGTCGTGGTTCCGCTGATTACTCCTCTCA–TAO/IBRQ-3.’

### In Vivo Efficacy Study In Human CCA and HCC Xenograft Models

To evaluate the *in vivo* therapeutic efficacy of oncolytic virus MORV and VSV in subcutaneous human cholangiocarcinoma (CCA) and hepatocellular carcinoma (HCC) xenograft models, 2 x 10^6^ HuCCT-1 (CCA) and Hep3B (HCC) cells (n= 7 mice per group) were subcutaneously inoculated into the right flanks of female athymic nude (NU/J) mice (Jackson laboratories, USA). When average tumors reached 100-200 mm^3^, mice were randomized into the respective study groups and dosed within 24 hours of randomization. Mice received two (HuCCT-1) or one (Hep3B) intra-tumoral (IT) injections, each one week apart with 50 µl containing PBS or 1×10^7^ TCID_50_ units of MORV or VSV. Conversely, mice in the Hep3B cohort received a single dose of PBS or 1 x 10^7^ TCID_50_ units of MORV or VSV. Tumor volume and body weight of the mice were monitored. Mice were euthanized when adverse effects were observed or when tumor size was larger than 2000 mm³. Tumor volume was calculated using the following equation: (longest diameter * shortest diameter^2^)/2. Tumor images were taken prior to resection, and tumor weight recorded post-resection.

### In Vivo Efficacy in Syngeneic, Orthotopic Mouse Model of Cholangiocarcinoma (CCA)

Fifty (50) female C57BL/6J mice (n= 10 mice per group) were purchased from the Jackson laboratories. Mice were anesthetized using 1.5-3% isoflurane. Under deep anesthesia, the abdominal cavity was opened by a 1 cm incision below the xiphoid process. A sterile cotton tipped applicator was used to expose the superolateral aspect of the medial lobe of the liver. Using a 27-gauge needle, 20 µL of standard media containing 0.75 x 10^6^ SB cells (murine CCA cells) ^34–36^ was injected into the lateral aspect of the medial lobe. A cotton tipped applicator was held over the injection site to prevent cell leakage and blood loss. Subsequently, the abdominal wall and skin were closed in separate layers with absorbable chromic 3–0 gut and nylon 4-0 skin suture material. All virus injections were initiated seven days post SB cells implantation. For the experiments employing oncolytic viruses, mice were randomly assigned to vehicle PBS, MORV 1 x 10^7^ TCID5_0_ or MORV 1 x 10^8^ TCID_50_, VSV 1 x 10^7^ TCID_50_ or VSV 1 x 10^8^ TCID_50_. Single intraperitoneal (IP) injections of viral preparation in 50µl was administered. Four (4) weeks following SB cell implantation, mice were sacrificed, and tumor, adjacent liver, spleen and blood were collected for downstream analysis. Animals were monitored daily for one week and then weekly for any changes in appearance or behavior, including any signs of morbidity or mortality. Images of resected tumors were taken at study termination.

### Analysis of Tumor-infiltrating Immune Cells

Upon excision, tumors from five mice per group were dissociated with gentleMACS^™^ Octo Dissociator (Miltenyi) according to the manufacturer’s protocol. CD45^+^ cells were isolated by CD45 (TIL) mouse microbeads (Miltenyi). Cells were incubated with Fixable Viability Stain 510 (BD Horizon™) for 15 minutes followed by anti-Fc blocking reagent (Miltenyi) for 10 minutes prior to surface staining. Cells were stained, followed by data acquisition on a Miltenyi MACSQuant® Analyzer 10 optical bench flow cytometer. All antibodies were used following the manufacturer’s recommendation. Fluorescence Minus One control was used for each independent experiment to establish gating. For intracellular staining of granzyme B, cells were stained using the intracellular staining kit (Miltenyi). Analysis was performed using FlowJo™ (TreeStar). Forward scatter (FSC) and side scatter (SSC) were used to exclude cell debris and doublets.

### Flow Cytometry Analysis Antibodies

The following antibodies were used for flow cytometry staining: F4/80-PE (REA126, Miltenyi), CD11b-PE-Cy5 (M1/70, eBioscience), CD206-PE-Cy7 (C068C2, BioLegend®), F4/80-PE-Vio®770 (REA126, Miltenyi), CD11c-APC (REA754, Miltenyi), Ly6G-PE (Rat 1A8, Miltenyi), Ly6C-APC-Vio®770 (REA796, Miltenyi) CD3-APC-Vio®770 (REA641, Miltenyi), CD8-BV421 (53-6.7, BD Horizon™), CD11a-PE-Vio®770 (REA880, Miltenyi), PD-1-PerCP-Vio®700 (REA802, Miltenyi), granzyme B-PE (REA226, Miltenyi).

### Murine Immunoglobulin Isotyping Following Treatment with MORV and VSV

For analysis of antibody production, serum samples from five mice in each group were obtained from whole blood collected in BD Microtainer tubes. Serum immunoglobulin subclass antibody responses (IgG1, IgGA, IgG2a, IgG2b, IgG3 and IgM) in mice were determined after infection with MORV and VSV using mouse Isotyping multiplex assay (MGAMMAG-300K, Millipore Sigma, USA) by Luminex xMAP technology as recommended by the manufacture.

### Serum Cytokines

Mouse IFN-α & IFN-β 2-Plex Mouse Panel (EPX02A-22187-901, ThermoFischer, USA) was used to measure serum levels of IFN-α and IFN-β. Blood chemistry (serum albumin, blood urea nitrogen, serum cholesterol) analysis was performed in an Abaxis Piccolo Xpress chemistry analyzer (Abaxis, Union City, CA, USA),

### Histopathological Analysis

Histopathological evaluation on H&E staining images for assessment of any abnormal changes in brain, liver and spleen were reviewed by a board-certified pathologist (Dr. Naomi Gades, Mayo Clinic in Arizona). HALO v3.1.1076.379 was used to measure the percentage tumor necrotic area.

### TUNEL Assay Immunohistochemistry

Liver histology was performed using tissue fixed in 10% formalin, dehydrated, and embedded in paraffin. Terminal deoxynucleotidyl transferase deoxyuridine triphosphate nick-end labeling (TUNEL) assay was carried out using the ApopTag Peroxidase In Situ Apoptosis Detection Kit (Millipore Sigma, St. Louis, MO); diaminobenzidine (DAB) was used as a peroxidase substrate (Vector Laboratories, Burlingame, CA); 0.5% methyl green was used for the counterstain as previously described.^58^ Positive cells were determined in five representative fields of adjacent liver and five usual fields in tumor for each group.

### Statistical Analysis

All values were expressed as the mean±SD (standard deviation), and the results were analyzed by one-way analysis of variance followed by the Tukey test for multiple comparisons and the Kaplan-Meier method for survival, using statistical software in GraphPad Prism, version 8 (GraphPad Software, Inc, La Jolla, California, USA). A *P* value less than .05 was considered significant.

## Acknowledgments

We thank the personnel of the Animal Facility and Histology Core at Mayo Clinic in Scottsdale, Arizona, for their assistance during the animal studies, including characterization of the toxicologic, pharmacologic, and efficacy effects of MORV and VSV in animal models.

## Author Contributions

BMN and MJB contributed to study concept and design, data analysis, interpretation of data, and drafting of the manuscript. DGD and SI contributed to reagents. SI, EL, ATB, YZ, NM, JP, FA, CD, OB, PLSUJ, and MA contributed to acquisition of data, interpretation of data, and critical revision of the manuscript. BJ, KB, MAB, DGD, LRR, RV, JMB, MTB, SRP and MJC contributed to drafting and critical revision of the manuscript. All authors approved the final, submitted version of the manuscript.

## Abbreviations

ALP: alkaline phosphatase
ALT: alanine aminotransferase
Anti-VSV-G: anti-vesicular stomatitis virus glycoprotein antibody
AST: aspartate aminotransferase
ATCC: American Tissue Culture Collection
CBC: complete blood count
CCA: cholangiocarcinoma
CPE: cytopathic effect
CTLs: cytotoxic T lymphocytes
DMEM: Dulbecco’s Modified Eagle Medium
DPBS: Dulbecco’s Phosphate-buffered Saline
ELISA: enzyme-linked immunosorbent assay
G-MDSCs: granulocytic myeloid-derived suppressor cells
GzmB: granzyme B
h: human
H&E: hematoxylin and eosin
HCC: hepatocellular carcinoma
HGB: hemoglobin
IFN: interferon
IFN-α: interferon-alpha
IFN-β: interferon-beta
IgG1: immunoglobulin G1
IgG2a: immunoglobulin G2a
IgG2b: immunoglobulin G2b
IgG3: immunoglobulin G3
IgGA: immunoglobulin A
IgM: immunoglobulin M
LDLR KO: HAP-1 LDLR knock-out cells
LDLR: low-density lipoprotein receptor
m: mouse
M-MDSCs: mononuclear myeloid-derived suppressor cells
MOI: multiplicity of infection
MORV: Morreton virus
MTS: 3-(4,5-dimethylthiazol-2-yl)-5-(3-carboxymethoxyphenyl)-2-(4 sulfophenyl) 2H-tetrazolium
PBS: phosphate buffered saline
PD-1: programmed cell death protein 1
PE: phycoerythrin
PT13-D22: patient 13 treated with VSV-hIFN-β day 22 sample
PT14-22: patient 14 treated with VSV-hIFN-β, day 22 sample
qRT-PCR: quantitative real-time polymerase chain reaction
RBC: red blood cells
RNA: ribonucleic acid
RPMI 1640: Roswell Park Memorial Institute cell culture medium
RWA: red cell distribution width
RT: Room temperature
TAMs: tumor associated macrophages
TCID_50_: 50% tissue culture infectious dose
Vero: African green monkey kidney cell line
VSV: vesicular stomatitis virus
VSV-hIFN-β: vesicular stomatitis virus expressing human interferon-beta
WT: wild type

## SUPPLEMENTARY FIGURES AND TABLES

### Legends

**Supplementary Figure 1.** <u>Glycoprotein Based Phylogeny of MORV</u>. Phylogenic tree based on MORV glycoprotein and others close, yet distinct *Vesiculoviruses* using the Maximum Likelihood method and JTT matrix-based model.

**Supplementary Figure 2.** <u>MORV and VSV Sensitivity to Human Type I Interferon (IFN)</u>. **A,** Phase-contrast microscopic images of A549 cells infected treated with human interferon-alpha (IFN-α) and infected with MORV and VSV at a multiplicity of infection (MOI) of 0.01 for 48 hours post infection. **B,** A panel of human tumor cells (2×10^4^) cells were infected with MORV and VSV at MOI of 5. We individually measured interferon-beta levels (IFN-β) 18 hours after infection in the supernatants of infected cells. The average of three independent experiments are plotted.

**Supplementary Figure 3.** <u>Intranasal Administration of MORV and VSV Are Well Tolerated in Laboratory Mice</u>. **A,** Changes in the blood parameters, including monocyte, granulocyte, red cell distribution width (RWA), hemoglobin (HGB), red blood cells (RBC) and platelets after intranasal treatments with MORV and VSV. **B,** Survival plots of mice treated with increasing intranasal doses of MORV and VSV (1×10^7^ TCID_50_/kg, 1×10^8^ TCID_50_/kg, 1×10^9^ TCID_50_/kg and 1×10^10^ TCID_50_/kg).

**Supplementary Figure 4.** <u>Quantitation of Viral N Gene mRNA in Mouse Tissues Following MORV and VSV Infection.</u> Quantification of viral nucleoprotein gene (MORV-N and VSV-N) in the brain, blood, liver, and spleen of immunocompetent mice treated with intranasal doses of MORV and VSV.

**Supplementary Figure 5.** <u>Individual Tumor Volume and Body weight (Tumor Xenografts</u>). Graphs showing the effect of treatment with single or multiple doses of MORV and VSV on individual body weight for HuCCT-1 (A) and Hep3B (B).

**Supplementary Figure 6.** <u>Real-Time Analysis of MORV-Induced Apoptosis</u> *<u>in vitro</u>* <u>and Quantification of Apoptotic Tumor Cells From *In vivo* Studies</u>**. A,** Murine cholangiocarcinoma (SB) cells (1 x10^4^ per well) were infected with MORV and VSV at different MOIs (10, 1, 0.1, and 0.01), and apoptosis was measured every 6 hours for 48 hours using Annexin V Red in the IncuCyte S3 system (Essen Bioscience, USA). Images were taken at 48 hours post-infection (MOI= 10). Data were plotted as mean ± SEM. **B,** graphs of terminal deoxynucleotidyl transferase dUTP nick-end labeling (TUNEL) assay on syngeneic tumor model of CCA.

**Supplementary Figure 7.** <u>Tumor Nodules, Liver Function Tests and Expression of MORV and VSV Genes In Orthotopic Murine CCA Model.</u> **A,** Representative image of tumor nodules in the control group (PBS) compared to MORV and VSV groups. **B,** Analysis of the expression of type I interferon (INF-α and INF-β), biochemical parameters (Bilirubin, albumin, cholesterol, and blood urea nitrogen) and **C**, quantification of viral nucleoprotein gene (MORV-N and VSV-N) in the tumor nodules after four weeks of treatment with MORV and VSV.

**Supplementary Figure 8.** <u>MORV Promotes Higher CTLs Infiltration in Murine CCA Tumor Microenvironment</u>. Immune profiling of PBS and viruses (MORV and VSV) treated syngeneic mice showing percentage of tumor infiltrating immune cells, including tumor-associated macrophages (TAMs), myeloid-derived suppressor cells (M-MDSCs), granulocytic myeloid-derived suppressor cells (G-MDSCs), programmed cell death protein 1 (PD1+), granzyme (GzmB+) reactive CTLs and natural killer (NK) cells. Unpaired T test and ordinary one-way ANOVA were used to establish significance (*P*=0.05).

**Supplementary Figure 9.** <u>Analysis of Virus-specific Immunoglobulin Levels of MORV Compared to VSV</u>. Analysis of the immunoglobulin profile of syngeneic mice treated with intraperitoneal injection of PBS or 1×10^7^ or 1×10^8^ TCID_50_ units of MORV or VSV. Unpaired T test and ordinary one-way ANOVA were used to establish significance (*P*=0.05).

**Supplementary Figure 10.** <u>Changes in Mouse Weight, Spleen Weight and Metastatic Nodules Following Treatment With MORV and VSV</u>. Changes in mouse weight, spleen weight and metastatic nodules following treatment with MORV and VSV. Evaluation of changes in body weight (**A**), spleen weight (**B**), and tumor metastasis in the intestine (**C**) of mice treated with MORV and VSV.

**Supplementary Table 1.** <u>List of Glycoproteins of known</u> *<u>Vesiculoviruses</u>* <u>Genetically Close to VSV.</u> List of Glycoproteins used to generate the Phylogeny Tree.

**Supplementary Table 2.** <u>Liver Cancer Cell lines</u>. A complete list of tumor and normal human and murine cell lines used, including media requirements and source are compiled in this table.

**Supplementary Table 3**. <u>Survival of Mice Treated with Intranasal Doses of MORV and VSV (45 days)</u>.

