## Supplementary Figures for "Characterization of Morreton Virus (MORV) as a Novel Oncolytic Virotherapy Platform for Liver Cancers"

### Supplementary  Figures and Tables

**Supplementary Figure 1.** Glycoprotein based Phylogeny of Morreton Virus

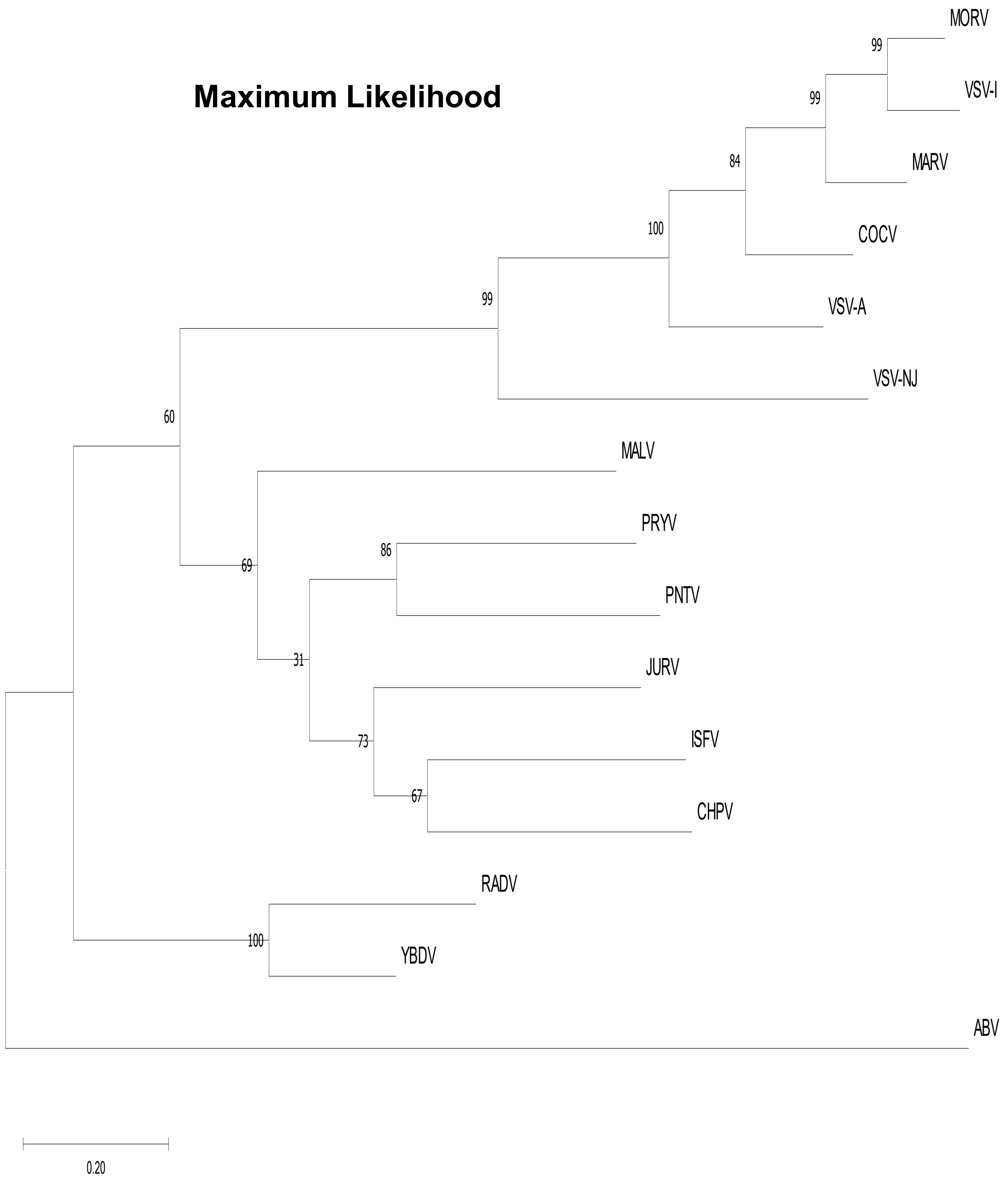

**Supplementary Figure 2**: MORV and VSV sensitivity to human type I interferon (IFN)

A

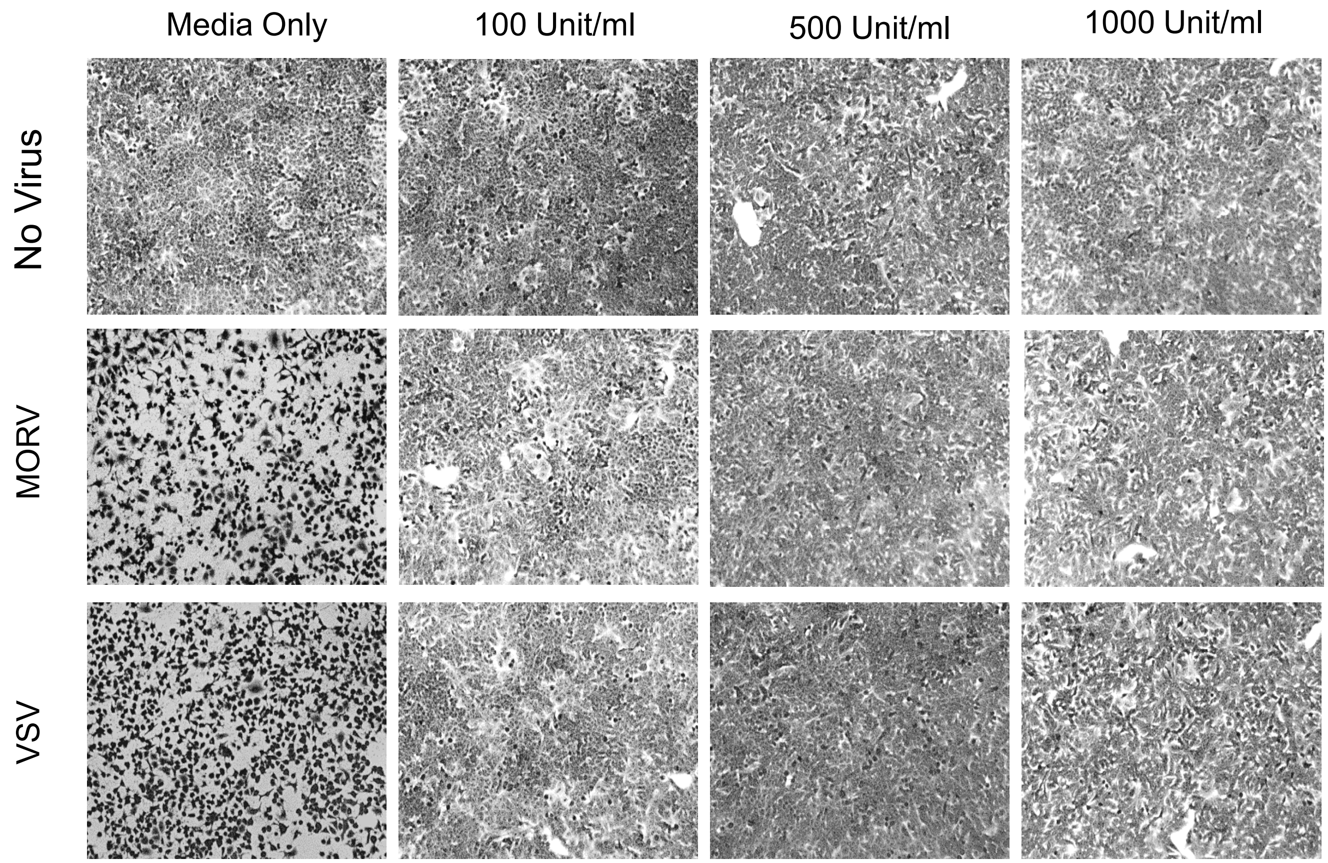


B

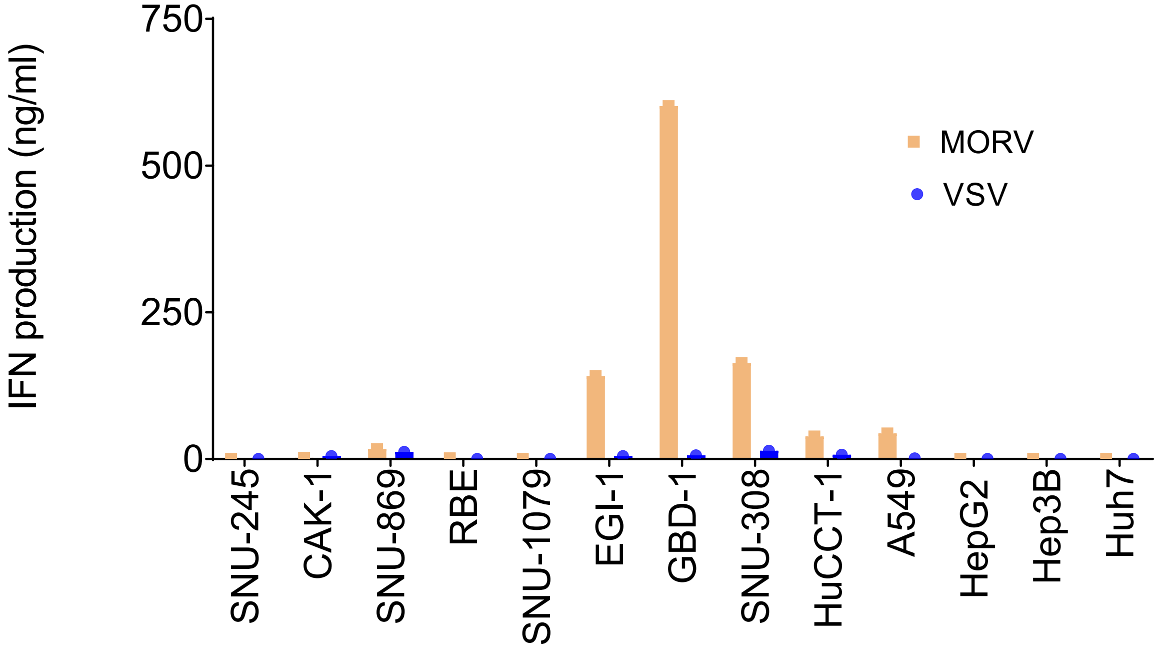

**Supplementary Figure 3.** Intranasal administration of MORV and VSV are  well tolerated in Laboratory Mice

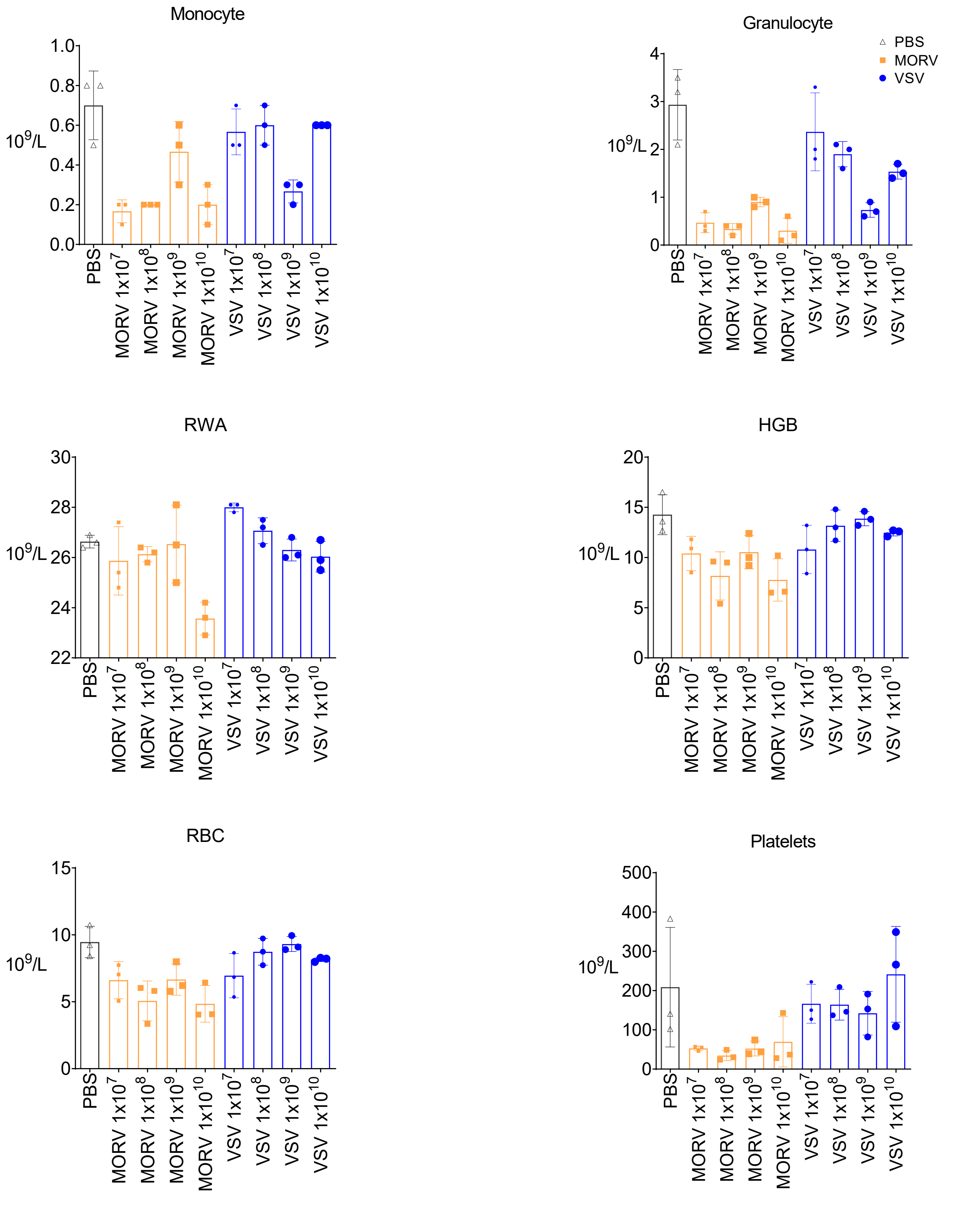

**Supplementary Figure 4:** Quantitation of viral N gene mRNA in mouse tissues following MORV and VSV infection

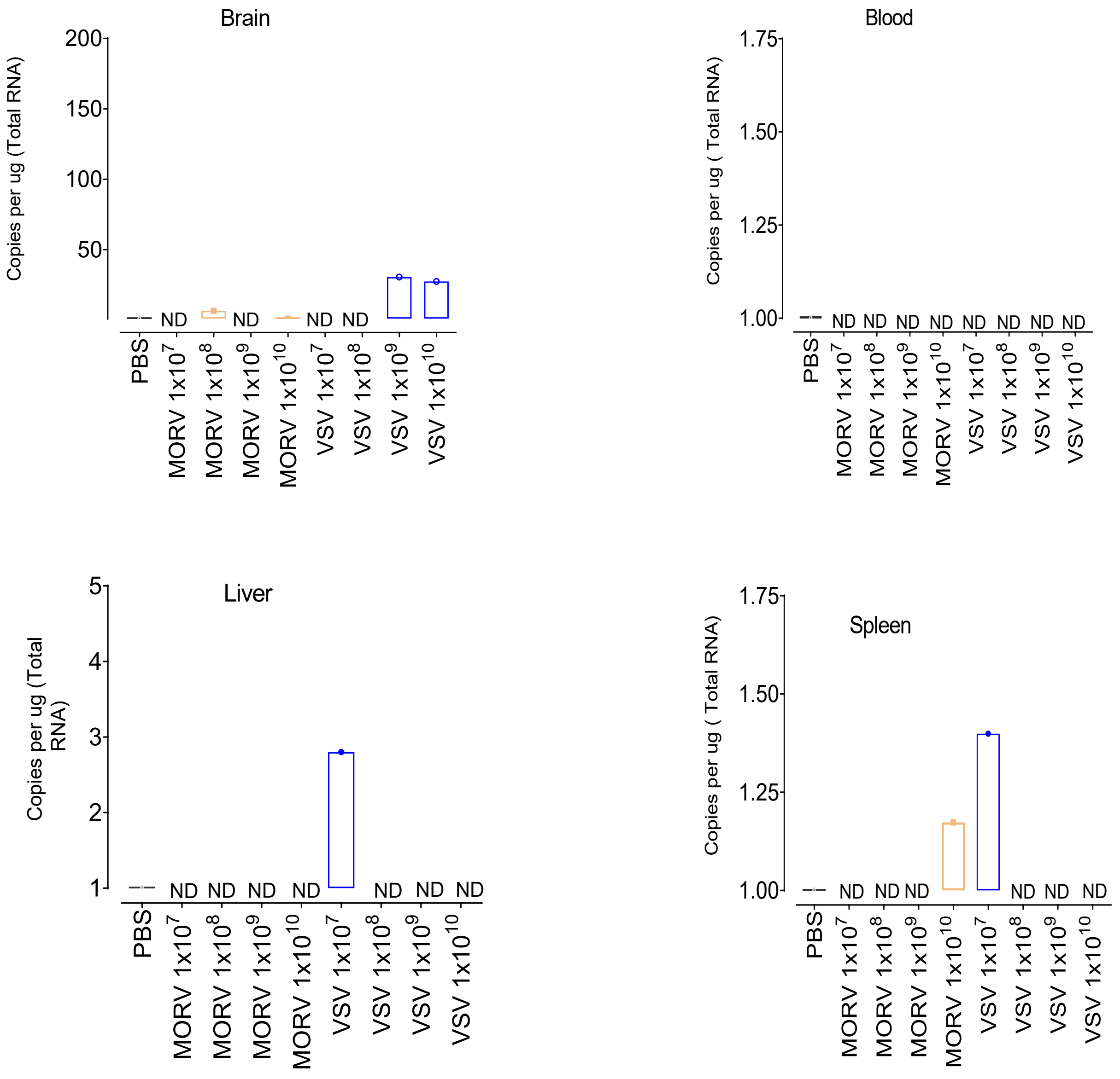


***Dosage:** TCID50/kg;* ***ND**: Non detected or < 1000 copies per ng of total RNA*

**Supplementary Figure 5:** Individual tumor volume and Body weight ( tumor xenografts)

A

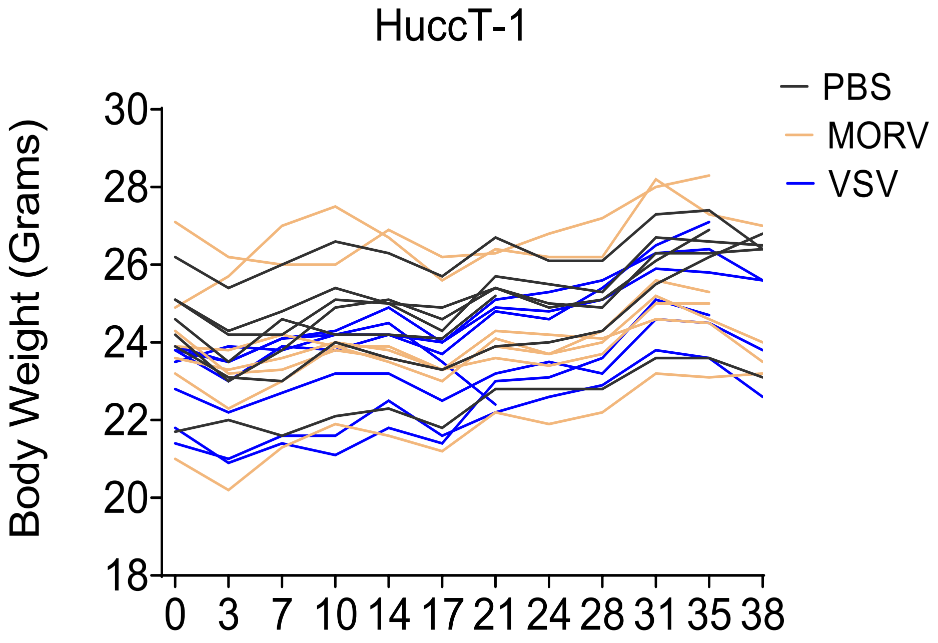


B

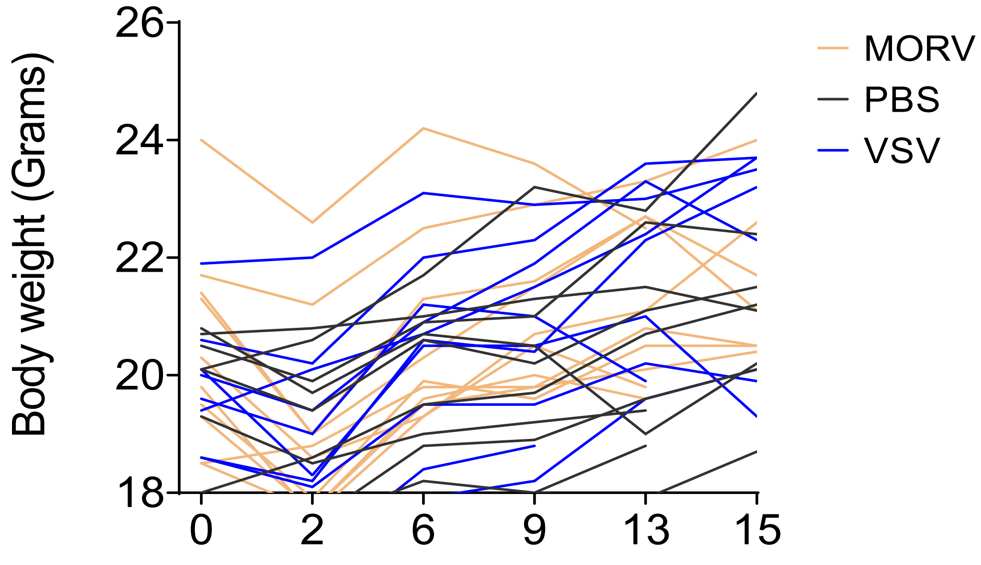

**Supplementary Figure 6.** Real-time analysis of MORV-induced apoptosis *in vitro* and quantification of apoptotic cells from *in vivo*

A

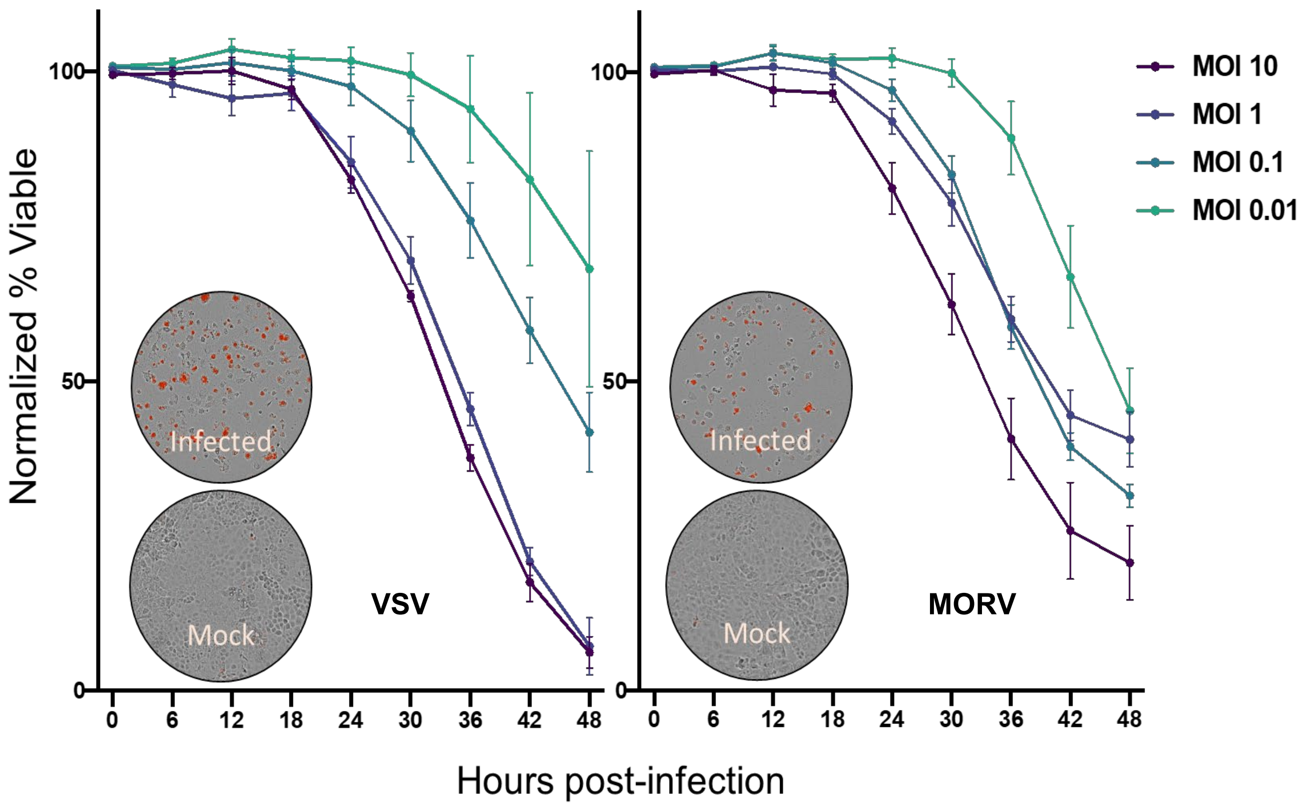


B

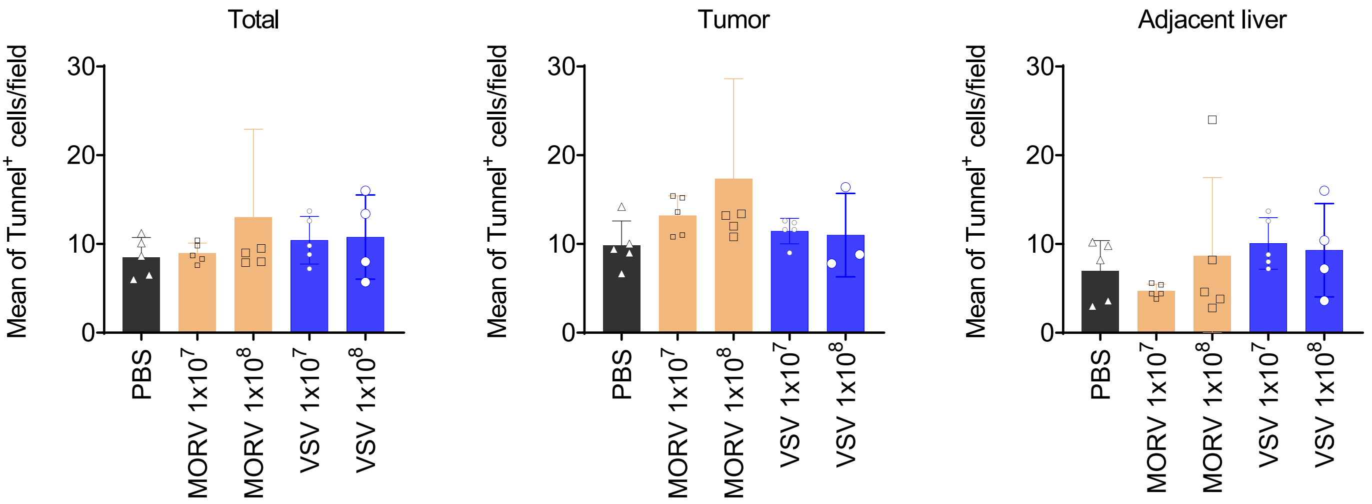

**Supplementary Figure 7.** Tumor nodules, liver function tests and expression of MORV and VSV genes in orthotopic murine model of CCA

A

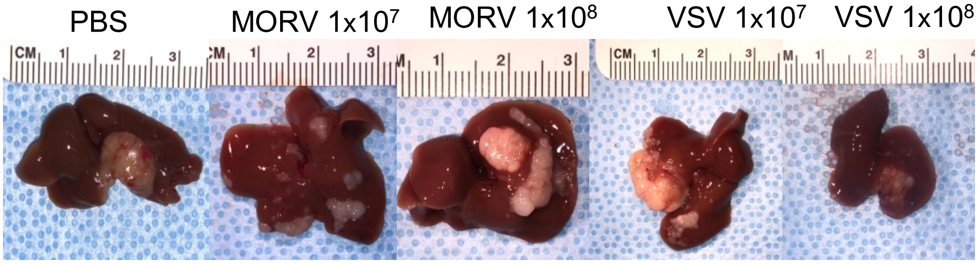


B

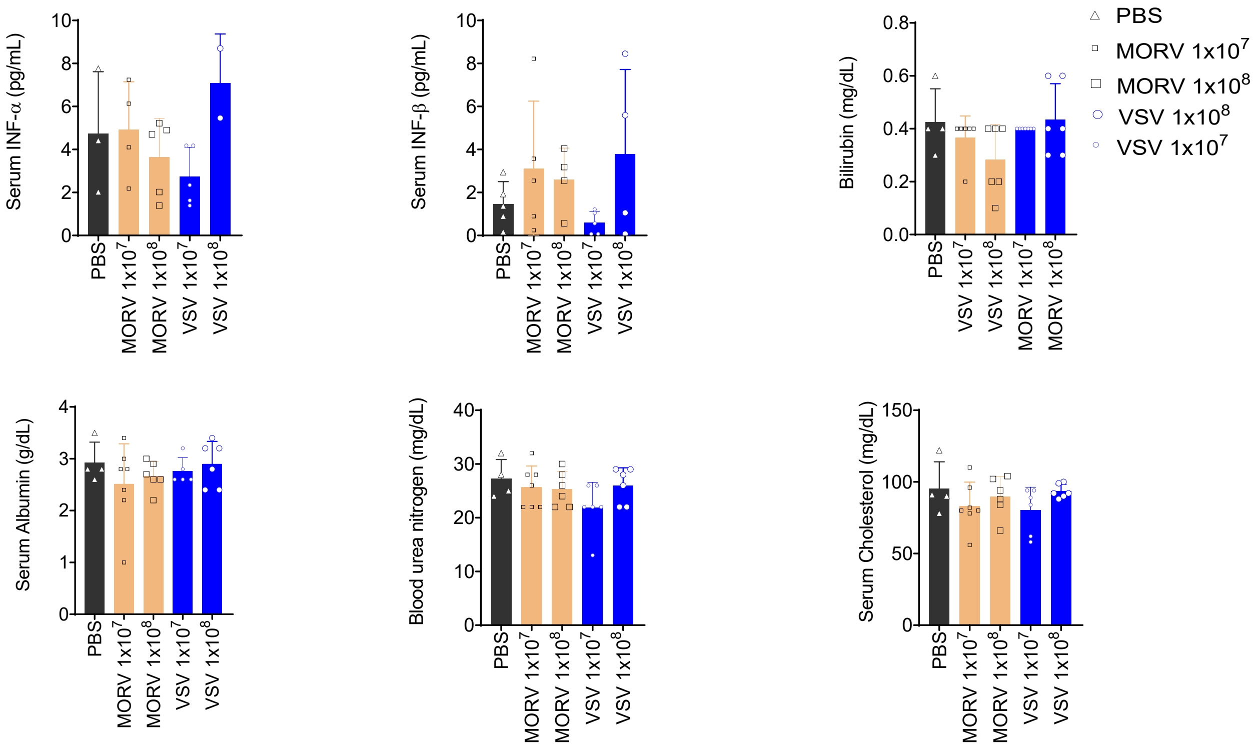


C

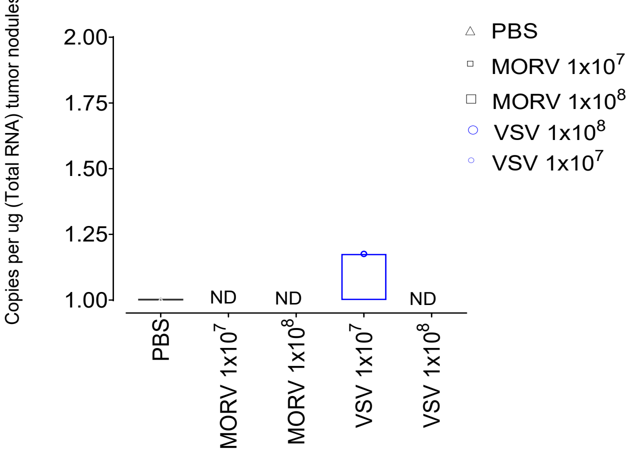

**Supplementary Figure 8.** MORV promote a higher CTLs infiltration in Murine CCA tumor microenvironment

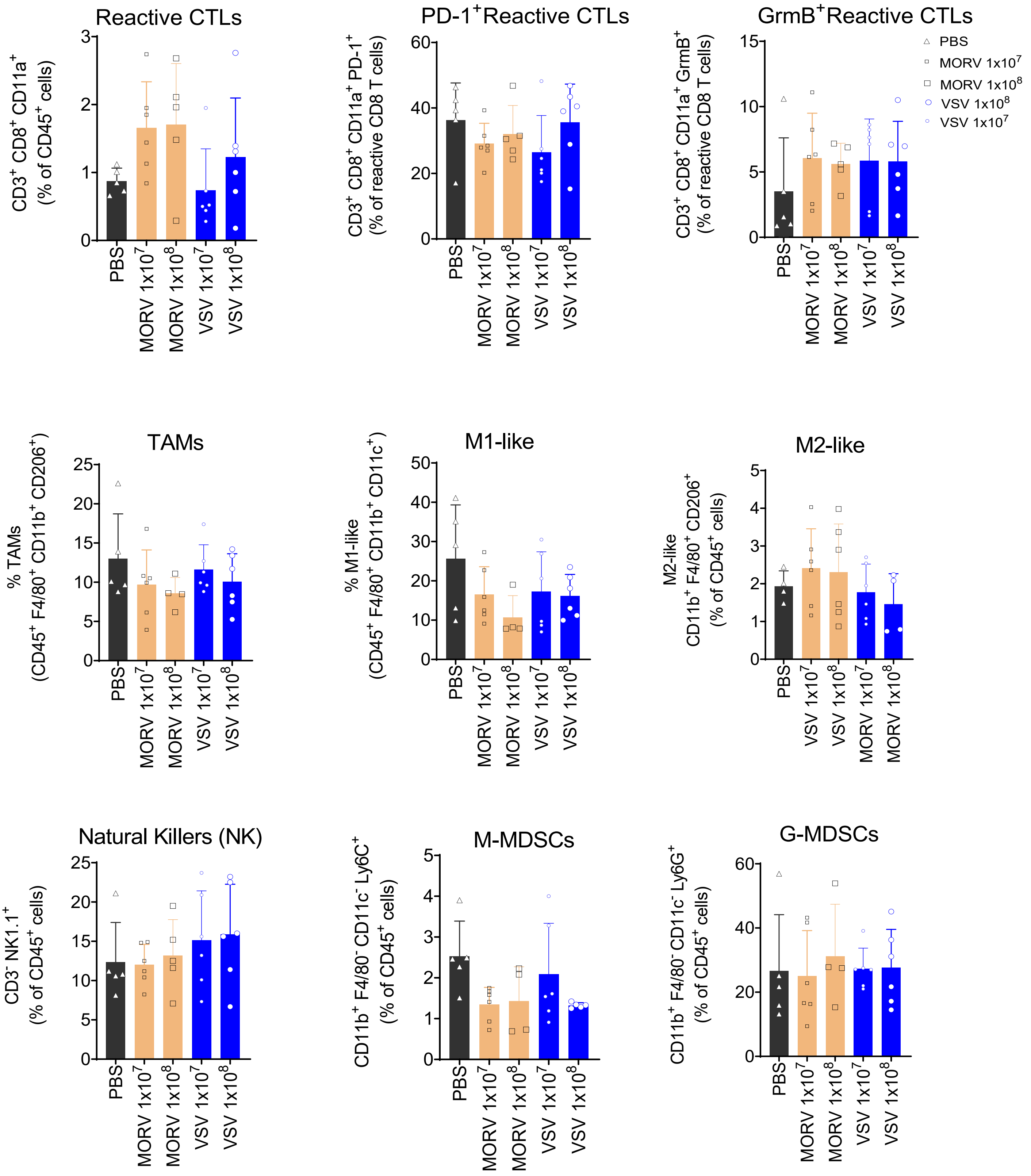

**Supplementary Figure 9.** Viral specific Immunoglobulin levels of MORV compared to VSV

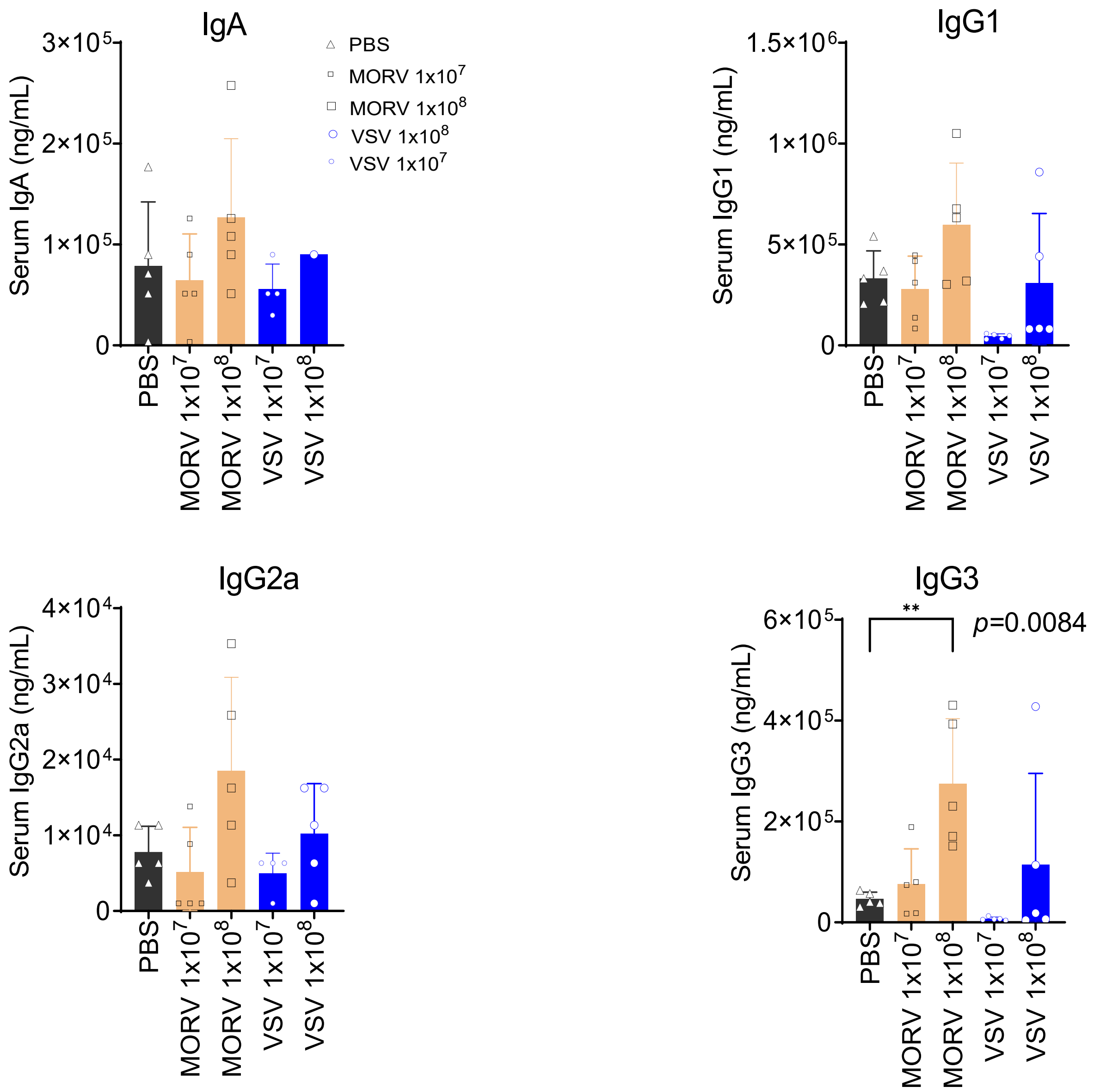

**Supplementary Figure 10.** Changes in mouse weight, spleen weight and metastatic nodules following treatment with MORV and VSV

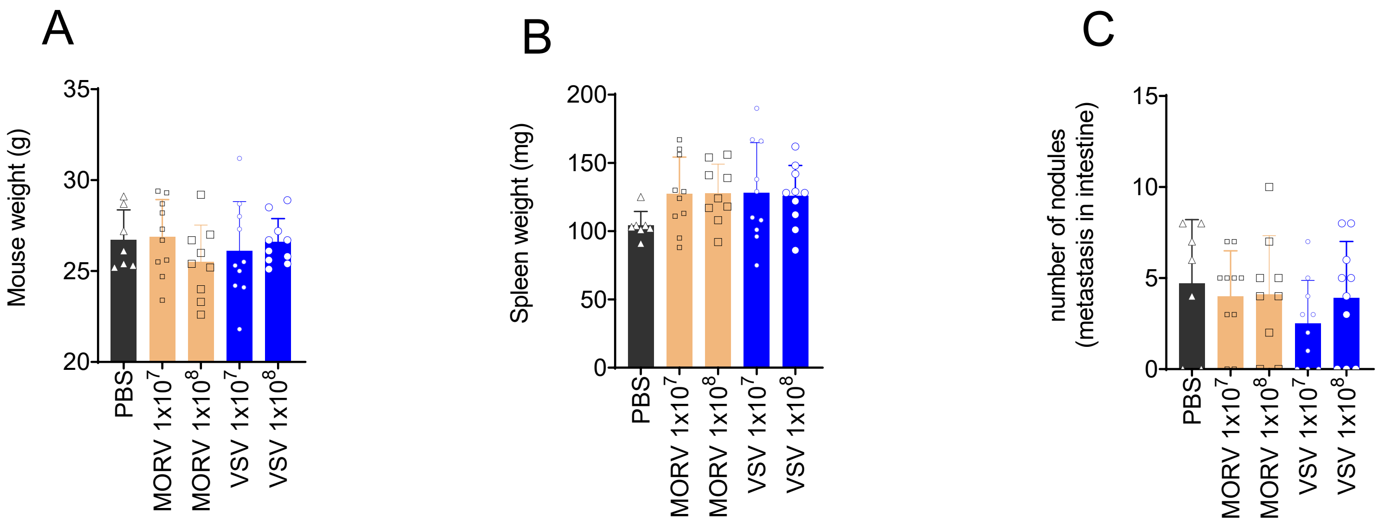

**Supplementary Table 1.** List of Glycoproteins of know Vesiculoviruses genetically close to VSV.

| Protein Reference sequence | Glycoprotein (G) | Strain |
|---|---|---|
| >NP_041715.1 | Glycoprotein (G protein) precursor [Vesicular stomatitis Indiana virus] | >VSV-I |
| >YP_009362085.1 | Glycoprotein [Morreton vesiculovirus] | >MORV |
| >YP_009091829.1 | Glycoprotein [Maraba virus] | >MARV |
| >AAC02712.1 | Glycoprotein G [Cocal virus] | >COCV |
| >ACB47442.1 | Glycoprotein [Vesicular stomatitis Alagoas virus] | >VSV-A |
| >YP_009047084.1 | Glycoprotein [Vesicular stomatitis New Jersey virus] | >VSV-NJ |
| >YP_007641385.1 | Glycoprotein [Isfahan virus] | >ISFV |
| >QNS83659.1 | Glycoprotein [Chandipura virus] | >CHPV |
| >YP_009505535.1 | Glycoprotein [Piry virus] | >PRYV |
| >AEG25354.1 | Glycoprotein [Perinet vesiculovirus] | >PNTV |
| >YP_009094177.1 | Glycoprotein [Malpais Spring vesiculovirus] | >MALV |
| >YP_009513006.1 | Glycoprotein [Jurona vesiculovirus] | >JURV |
| >YP_008767242.1 | Glycoprotein G [American bat vesiculovirus TFFN-2013] | >ABV |
| >YP_009505540.1 | Glycoprotein [Radi vesiculovirus] | >RADV |
| >YP_009094276.1 | Glycoprotein [Yug Bogdanovac vesiculovirus] | >YBDV |

**Supplementary Table 2.** Liver Cancer Cell lines

| Cell line | Site | Media | Source |
|---|---|---|---|
| CAK-1 | eCCA | RPMI1640 + 10% FBS + AA | Gift from Dr. Gores |
| EGI-1 | eCCA | RPMI1640 + 10% FBS + AA | DSMZ |
| GBD-1 | GBC/eCCA | RPMI1640 + 10% FBS + AA | Gift from Dr. Gores |
| H69 | Normal cholangiocyte | DMEM/F12 + 10% FBS +GFs | Gift from Dr. Gores |
| HuCCT1 | iCCA | DMEM + 10% FBS + AA | JCRB |
| PAX-42 | CCA | DMEM + 10% FBS + AA | Gift from Dr. Truty |
| RBE | iCCA | RPMI1640 + 10% FBS + AA | RIKEN |
| SNU-1079 | iCCA | RPMI1640 + 10% FBS + AA | KCLB |
| SNU-245 | eCCA | RPMI1640 + 10% FBS + AA | KCLB |
| SNU-308 | GBC/eCCA | RPMI1640 + 10% FBS + AA | KCLB |
| SNU-869 | Ampulla of Vater | RPMI1640 + 10% FBS + AA | KCLB |
| SB | Murine CCA | DMEM + 10% FBS + AA | Dr. Rizvi |
| Hep3B | HCC | DMEM + 10% FBS + AA | ATCC |
| HepG2 | HCC | DMEM + 10% FBS + AA | ATCC |
| Huh7 | HCC | DMEM + 10% FBS + AA | ATCC |
| Sk-Hep1 | HCC | DMEM + 10% FBS + AA | ATCC |
| R1LWT (clone 1 of RIL-175 cells) | Murine HCC | DMEM + 10% FBS + AA | Dr. Duda |
| R2LWT ( Clone 2 of RIL-175 cells) | Murine HCC | DMEM + 10% FBS + AA | Dr. Duda |
| HCA-1 | Murine HCC | DMEM + 10% FBS + AA | Dr. Duda |

**Supplementary Table 3.** Survival of Mice Treated with Intranasal Doses of MORV and VSV (45 days)

| Virus | 1 x $10^7$ TCID50/kg | 1 x $10^8$ TCID50/kg | 1 x $10^9$ TCID50/kg | 1 x $10^{10}$ TCID50/kg |
|---|---|---|---|---|
| **MORV** | 3/3 (100%) | 3/3 (100%) | 3/3 (100%) | 3/3 (100%) |
| **VSV** | 3/3 (100%) | 3/3 (100%) | 3/3 (100%) | 3/3 (100%) |
